# Competing calcium sensors orchestrate various patterns of synaptic transmission

**DOI:** 10.64898/2026.09.09.750532

**Authors:** Yinyun Li, Jules Lallouette, Iain Hepburn, Weiliang Chen, Erik De Schutter

## Abstract

Neurotransmission critically depends on both timing and efficacy, enabling neurons to encode information with highly accuracy in millisecond timescales. While synapses often express multiple calcium sensors, such as synaptotagmin-1 (syt1) and synaptotagmin-7 (syt7), the quantitative mechanisms by which these sensors regulate synchronous release (SR) and asynchronous release (AR) remain unresolved. We develop a biophysically detailed stochastic model of a presynaptic bouton that incorporates the distinct calcium-binding kinetics of syt1 and syt7 to dissect their roles in shaping synaptic release. We demonstrate how a rich repertoire of SR and AR patterns is influenced by calcium channel distribution, sensor quantity, the external calcium concentration and buffer properties. Importantly, the interplay between syt1 and syt7— through their distinct calcium affinities and exocytotic kinetics — constitutes a core mechanism for neural transmission. These findings establish calcium partitioning as a core mechanism driving the diverse release patterns of syt1 and syt7, accommodating even more complex multi-sensor environments.

## 1. Introduction

Synaptic vesicle release kinetics play a central role in neural signal transmission. The timing and efficiency of presynaptic vesicle fusion critically shape neural computation, short-term plasticity, and brain function (Abbott & Regehr, 2004; Kaeser & Regehr, 2014; Nicoll & Schmitz, 2005; Regehr, 2012; Südhof, 2013). In central synapse, multiple calcium sensors contribute to this process, notably synaptotagmin-1 (syt1) and synaptotagmin-7 (syt7)(Huson & Regehr, 2020; Goda & Stevens, 1994; Kaeser & Regehr, 2014; Sugita et al., 2002; Turecek & Regehr, 2018). Syt1 acts as a fast calcium sensor that mediates synchronous release (SR) (Bouazza-Arostegui et al., 2022; Geppert et al., 1994; Li et al., 1995; Nishiki & Augustine, 2004; Shin et al., 2009; Tagliatti et al., 2020); whereas syt7 has been proposed to function as a slower sensor underlying asynchronous release (AR) (Atluri & Regehr, 1998; Bacaj et al., 2013; Bose et al., 2024; Fukuda et al., 2004; Hui et al., 2005; Turecek & Regehr, 2018; Weingarten et al., 2024; Wen et al., 2010). Despite extensive characterization of these sensors individually, it remains unclear how their distinct biophysical properties are coordinated within the vesicle release machinery to generate different release modes.

Accumulating evidence suggests that syt1 and syt7 engage in both competitive and cooperative interactions within the release machinery. For example, syt1 knockout facilitates syt7-mediated release (Maximov & Südhof, 2005; Yoshihara & Littleton, 2002), and fast synaptotagmin isoforms regulate the relative contributions of SR and AR (Turecek & Regehr, 2019). In addition, high-frequency firing selectively recruits AR at mossy fiber boutons (MFB) (Chamberland et al., 2020), while syt7-mediated AR can enhance high-fidelity synchronous transmission at central synapses (Luo & Südhof, 2017). Together, these findings underscore the importance for a clear mechanistic characterization of syt7 to distinguish its function from syt1 and to allow deciphering of their complex interplay in shaping synaptic transmission.

At the molecular level, the C2A and C2B domains of syt7 bind calcium with distinct intrinsic affinities and kinetics (Brandt et al., 2012; Davis et al., 1999; Fukuda et al., 2004; Sugita et al., 2002; Voleti et al., 2017), providing insight into its functional differences from syt1. While calcium-binding kinetics of syt1 are well characterized (Davis et al., 1999; Geppert et al., 1994; Hui et al., 2005; Li et al., 1995; Nishiki & Augustine, 2004), for syt7 calcium binding kinetics were reported to be slow and its membrane-binding properties may explain the key role of its c2A domain in AR (Rao et al., 2017; Sugita et al., 2001, 2002; Vevea et.al., 2021; Volynski & Krishnakumar, 2018). A key factor that determines each syt’s function in neurotransmission in a shared synapse is which syt has a higher affinity for calcium. In addition, because each syt displays different releasing rates with fully bound calcium differential patterns of release kinetics are expected (Hui et al., 2005; Turecek & Regehr, 2019).

In parallel, the endogenous calcium buffer calmodulin (CaM) plays an important role in regulating release kinetics through calcium binding (Faas et al., 2011; Lou et.al., 2005;Matveev et al., 2004; Neher 1998;Neher & Sakaba, 2008; Schneggenburger & Neher, 2000, 2005; Timofeeva & Volynski, 2015; Li et al.,1995). CaM has been shown to shape vesicle release and short-term synaptic plasticity (STP) (Bollmann et al., 2000; Fogelson & Zucker, 1985; Neher & Sakaba, 2008; Timofeeva & Volynski, 2015). Because CaM promotes slow decays of calcium dynamics, it can gate long-lasting AR at the hippocampal mossy fiber to Ca3 pyramidal cell synapse (Chamberland et al., 2020). However, how CaM modulates syt1 and syt7 by competing with synaptotagmin sensors for binding calcium and shaping release dynamics remains unexplored.

Despite intensive investigation with advanced techniques of syt7 function in AR and its interaction with syt1 in different synaptic boutons across neuronal types, a generic and quantitative characterization of the mechanisms by which syt1 and syt7 coordinate in regulating AR and SR still require computational models and simulations for better understanding. STochastic Engine for Pathway Simulations (STEPS) provides a powerful platform that can simulate stochastic chemical reaction-diffusions in realistic 3-D morphologies at the nanoscale (Chen et al., 2022; Chen & De Schutter, 2017; Hepburn et al., 2012) together with vesicle release and recycling (Gallimore et al., 2025; Hepburn et al., 2024), with high performances in parallel simulations. However, syt7 ‘s function for AR was not included in the previous studies of a hippocampal en passant bouton (Gallimore et al., 2025).

In this work, we develop a detailed model of calcium-binding kinetics for syt7 and integrate it with syt1 in a simplified, half-hemispheric presynaptic mesh for STEPS simulation. By characterizing syt7 calcium binding kinetics, we simulate syt1 and syt7 triggered synaptic transmission with detailed vesicle release machinery. Our results show that differences in calcium-binding kinetics between syt1 and syt7, together with calcium buffering by CaM, give rise to a rich spectrum of SR and AR patterns across stimulation frequencies (10–100 Hz). By varying calcium channel distribution patterns, syt7 abundance, and extracellular calcium concentration, our model not only reproduces several key experimental observations in specific parameter regions, but also provides a broader platform for prediction. Our study reveals that partitioning of calcium by each sensor and buffer provides a core mechanism in regulating SR and AR.

## 2. Methods and Model

### 2.1 Properties of syt1 and syt7: intrinsic affinities to calcium and timescales

#### 2.1.1 Vesicle exocytosis by calcium binding to syts

Vesicle protein reactions leading to exocytosis are simulated in detail (Gallimore et al., 2025) and are summarized here. Vesicles are docked to active zones by binding of the vesicle surface molecule Rab3 to active zone proteins Rab3-interacting molecule (RIM) and Munc13 (M13). This complex later binds Munc18 (M18) and Syntaxin (SYX), as well as SNAP-25 and synaptobrevin to form a SNARE (Soluble N-ethylmaleimide-sensitive fusion protein (NSF) Attachemnt Receptor) (Grushin et al., 2019; Manca et al., 2019; Norman et al., 2023; Südhof & Rothman, 2009). The SNARE complex then binds synaptotagmin (syt) and complexin (CXN) to form SNARE_CXN_syt (Bera et al., 2022; Südhof & Rothman, 2009; Rothman et al.,2017; Zhou et al., 2015, 2017). In this model, we simplified these reactions: the Rab3 and RIM_M13 reaction is kept as a docking signal, but once vesicles are docked at the active zone, they will be equipped with SNARE_CXN_syt complex. Upon depolarization, calcium flows into the bouton and binds to syt in SNARE_CXN_syt complex, and eventually trigger membrane fusion and vesicle release (Ramakrishnan et al., 2018,2019,2020; Sun et al., 2007; Wu et al., 2022). Thus the final step of calcium binding to syt (Kaeser & Regehr, 2017; Regehr et al., 1994; Südhof, 2013) is critically important for the timing of signal transduction.

A schematic illustration (Fig.1) shows how a docked vesicle with SNARE_CXN_syt will be exocytosed when syt is fully bound by calcium that enters the bouton through voltage-gated calcium P-type channels. Initially each docking triangle in all active zones has one docked vesicle, while other vesicles are randomly distributed in the bouton (Fig. 1F). Upon calcium influx, only if syt binds calcium with both its C2A and C2B domains will the vesicle be exocytosed. The rate of exocytosis depends on how many SNARE_CXN_syt complexes with fully bound calcium are present on the vesicle: for 2 fully calcium bound SNARE_CXN_syt complexes the rate is *γ*_2_, for 3 the rate is 10 × *γ*_2_ and for 4 the rate is 100 × *γ*_2_. For syt1, *γ*_2_ = 3000*s*^−1^ (Gallimore et al., 2025); for syt7, *γ*_2_ = 10*s*^−1^ (Davis et al., 1999; Hui et al., 2005; Voleti et al., 2017). Calcium binding kinetics for C2A and C2B domains for both syt1 and syt7 are shown in Table 1.

**Figure 1.**
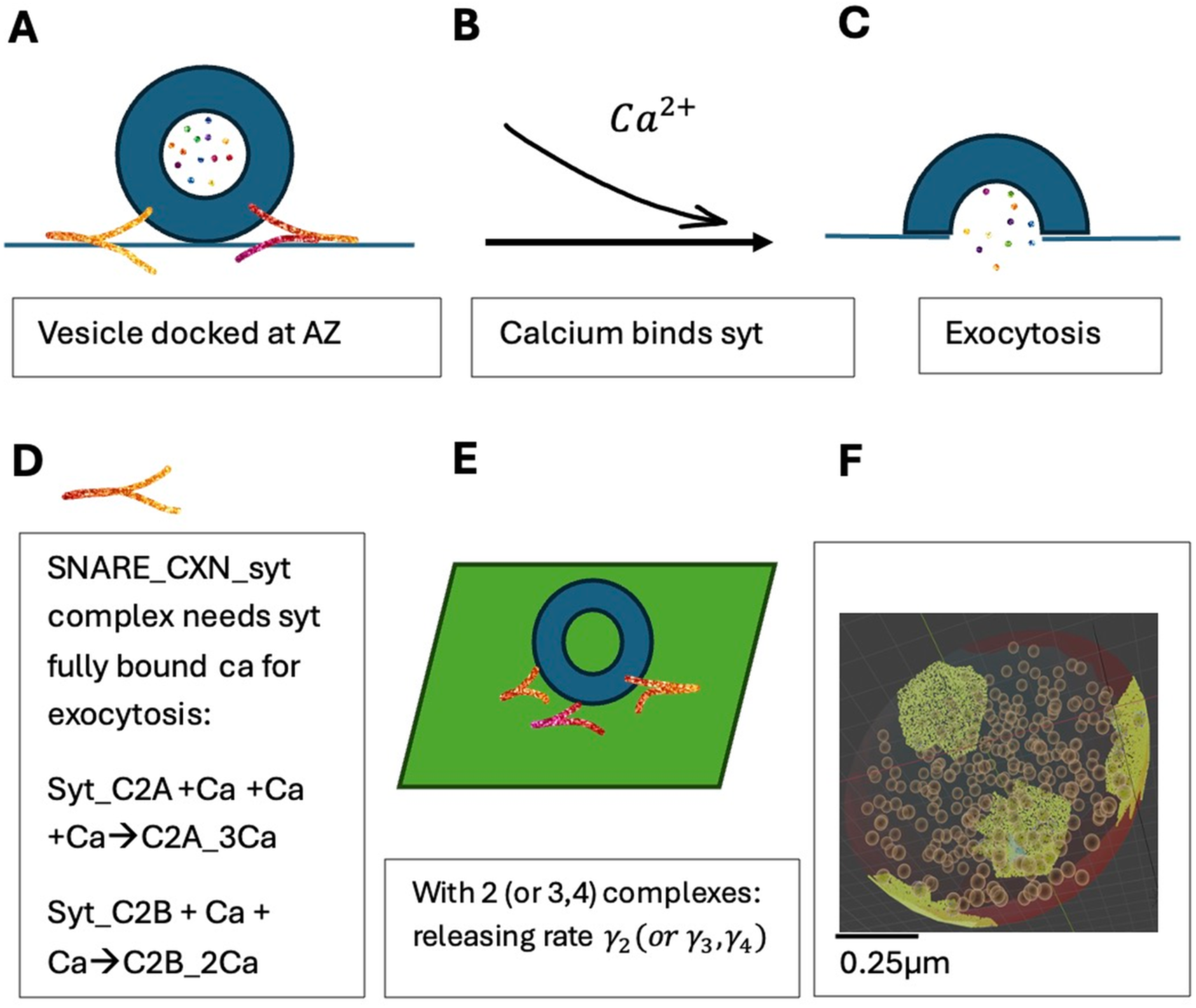
Schematic model of a docked vesicle with SNARE_CXN_syt complexes (A) and binding of calcium (B) for exocytosis (C). SNARE complexes need fully bound C2A and C2B domains to release (D) with rates of *γ*_2_, *γ*_3_, *γ*_4_ for 2, 3 or 4 SNARE_CXN_syt complexes (E). syt: synaptotagmin; the calcium binding model for C2A and C2B domains is identical for syt1 and syt7, with 3 Ca binding to C2A domain and 2 Ca binding to C2B domain, but with different kinetic rates (Table 1). (F) Blender plot of half-hemisphere mesh of a synaptic bouton showing the 4 active zones (yellow) and 300 vesicles.

**Table 1.** The calcium binding kinetic parameters for syt1 (Gallimore et al., 2025) and syt7 (our current model with best fit kinetic parameters), as well as CaM calcium binding to C-terminal and N-terminal (Gallimore et al., 2025).

| For syt1 | On rate[$M^{-1}s^{-1}$] | Off rate [$s^{-1}$] |
| --- | --- | --- |
| Syt1_C2A_1 | $k_1 = 2e9$ | $\kappa_1 = 1270$ |
| Syt1_C2A_2 | $k_2 = 2e9$ | $\kappa_2 = 227670$ |
| Syt1_C2A_3 | $k_3 = 2e9$ | $\kappa_3 = 12370$ |
| Syt1_C2B_1 | $f_1 = 2e9$ | $g_1 = 50000$ |
| Syt1_C2B_2 | $f_2 = 2e9$ | $g_2 = 25780$ |
| For syt7 |  |  |
| Syt7_C2A_1 | $k_1 = 9.18e7$ | $\kappa_1 = 58$ |
| Syt7_C2A_2 | $k_2 = 6.12e7$ | $\kappa_2 = 29$ |
| Syt7_C2A_3 | $k_3 = 3.06e7$ | $\kappa_3 = 10.875$ |
| Syt7_C2B_1 | $f_1 = 11.76e6$ | $g_1 = 65$ |
| Syt7_C2B_2 | $f_2 = 5.88e6$ | $g_2 = 32.5$ |
| For CaM |  |  |
| CaM_C1 | $16.8e07$ | $2.2e04$ |
| CaM_C2 | $2.5e07$ | $13$ |
| CaM_N1 | $15.4e08$ | $1.6e05$ |
| CaM_N2 | $3.2e10$ | $5.2e03$ |

#### 2.1.2 Time and spatial constants of calcium buffering

Following (Neher, 1998), we define a buffering time constant *τ* as:

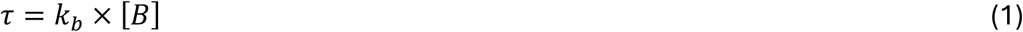

and a spatial buffering constant *λ* as:

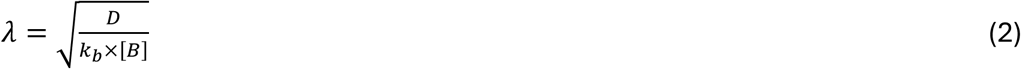

A stationary partition equation that can be used to describe the competition between buffers and sensors is derived in SI Section 4:

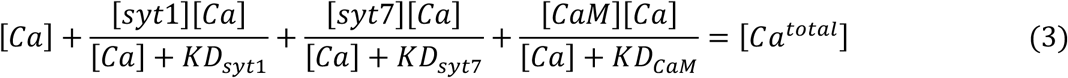

#### 2.1.3 Intrinsic and apparent affinities to calcium for syt1 and syt7

We derive an analytical expression for the intrinsic affinity to characterize the calcium binding properties for sensors syt1 and syt7. This framework can be applied to any calcium sensor or buffer which has similar calcium binding reactions.

The key result represents a detailed balance solution for the calcium binding kinetics for each sensor. The fully calcium bound state *S_fCa_* can be written as a function of kinetic rates, shown in eq. 4:

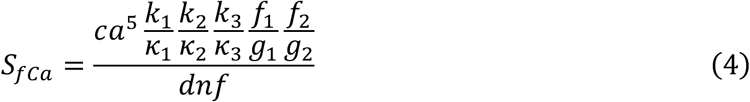

Where, *dnf* is a polynomial function shown in eq. 5:

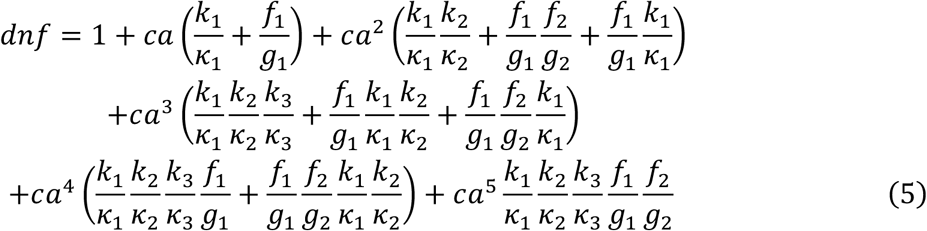

And *k*_1_, *k*_2_, *k*_3_*κ*_1_, *κ*_2_, *κ*_3_ are the binding and unbinding rates for C2A domain; *f*_1_, *f*_2_, *g*_1_, *g*_2_ are the binding rate and unbinding rate for the C2B domain (Table1).

The intrinsic affinity can be read out from a plot of eq. 4 as a function of calcium concentrations, i.e., for the y-axis at half maximum activation, the intrinsic affinity is the corresponding value on the x-axis. The same method can be applied to CaM, since CaM has a C-terminal and N terminal, each terminal binds 2 calcium ions, with kinetic rates from (Gallimore et al., 2025).

The apparent affinity observed experimentally depends, in addition, on calcium dependent lipid binding kinetics. This involves lipid binding on and off rates as well as lipid concentrations. How to compute the apparent affinity is described in SI Section 3. The apparent affinity is always lower than the intrinsic affinity.

#### 2.1.4 Timescale approximation for sensor binding calcium

Notice that the fully calcium bound state *S_fCa_* is a function of calcium concentration and ratios of unbinding and binding rates, but the timing information is missing. To investigate how fast each sensor can fully bind calcium, we have estimated the effective timescales for calcium binding kinetics.

The timescales of each sensor of binding kinetics can be analytically approximated by solving an effective 2-state model, where A is the state without calcium bound, and B is the state with fully calcium bound:

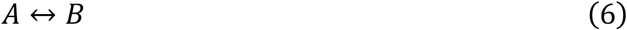

And the kinetic equation can be written as:

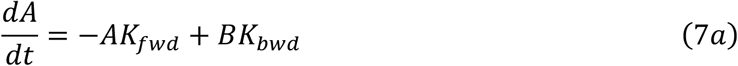

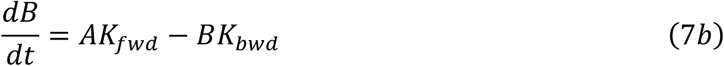

The effective forward rate *K_f_*_w*d*_ and backward rate *K_b_*_w*d*_ can be estimated by “*Rule of thumb*”(Gilbert, 1977), and expressed in the following equation:

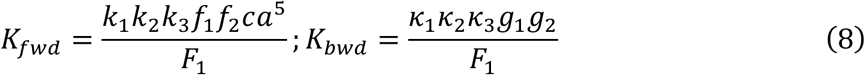

where,

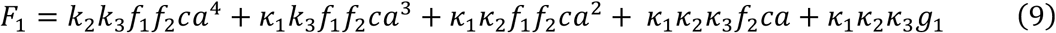

Notice that both *K_f_*_w*d*_ with unit of (*s*^−1^) and *K_b_*_w*d*_ with unit of (*s*^−1^) depend on calcium concentration.

The solution for eq. 7 has two eigenvalues, which can be written as:

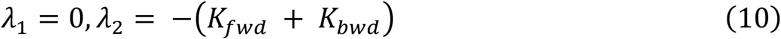

The timescales for the calcium binding kinetics can then be expressed as the inverse of the eigenvalue as:

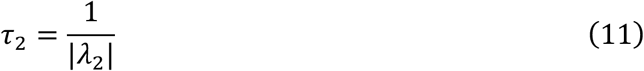

The effective time scale for CaM can be calculated by the same method. For other calcium buffers, parvalbumin (PV), calbindin (CB) and calretinin (CRTT), each bind 2 calcium ions with kinetics and concentrations adopted from (Gallimore et al., 2025). Their dissociation constants (KD) are at least ten times lower than CaM, indicating much higher affinity (Fig. 2- figure supplement 1) that will not directly compete with syt1 or syt7. Therefore, in this study we only illustrate the effect of CaM on syt1 and syt7 in causing different patterns of vesicle release.

**Figure 2.**
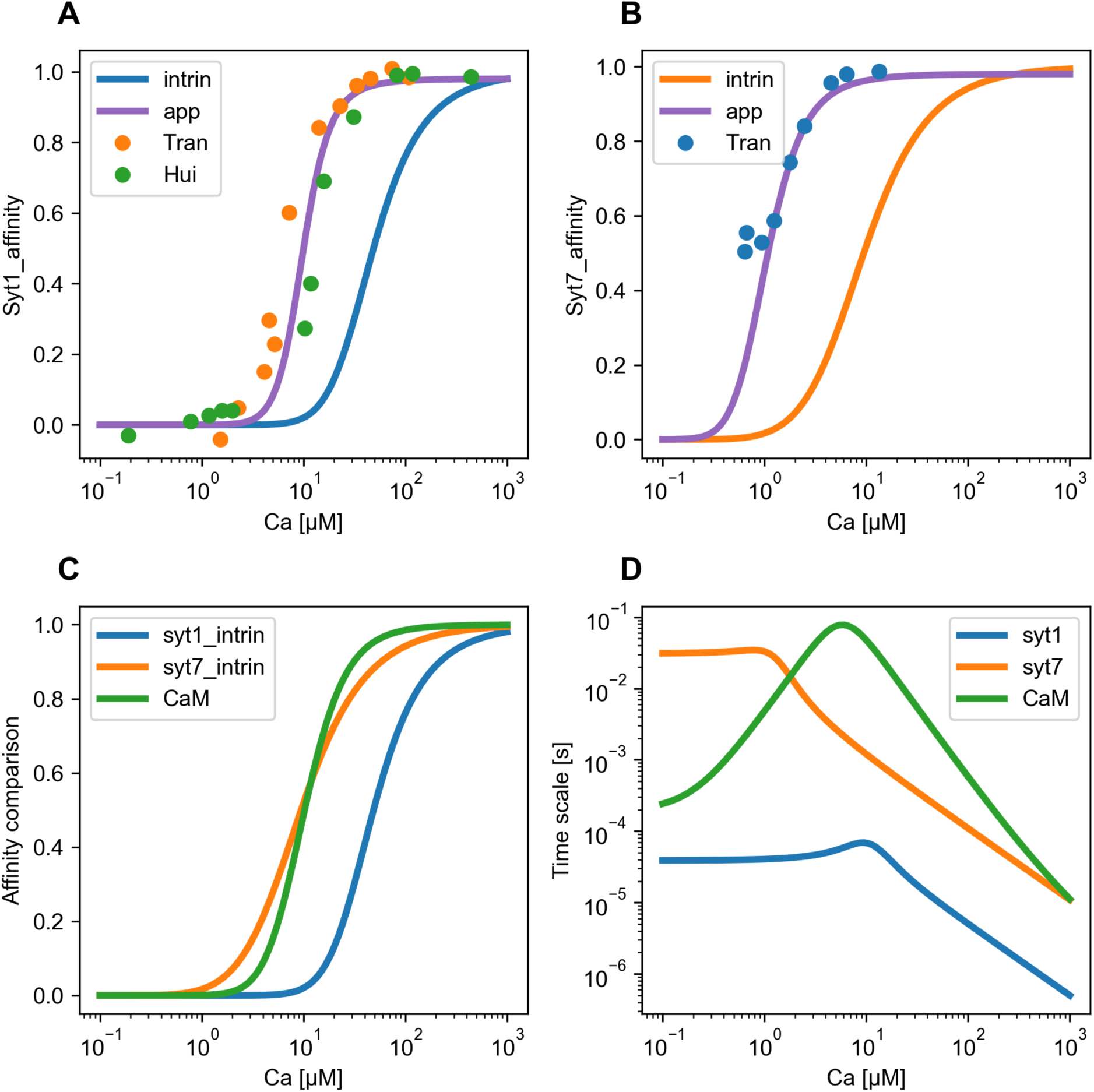
Comparison of intrinsic and apparent affinity between syt1 and syt7 and effective timescales for calcium binding kinetics. (A) apparent (app) and intrinsic (intrin) affinities for syt1. The dotted points are experimental data for calcium dependent lipid binding (Hui et al., 2005; Tran et al., 2019). (B) apparent and intrinsic affinities for syt7, compared to experimental observations. (C) intrinsic affinities of syt1, syt7, and CaM. (D) effective calcium binding timescales for syt1, syt7 and CaM.

### 2.2 STEPS simulations

We simulate the vesicle release kinetics with the STEPS software (Chen et al., 2022; Chen & De Schutter, 2017; Gillespie 1977;Hepburn et al., 2012, 2024). We constructed a half hemispheric mesh to mimic a presynaptic bouton, with radius 0.5 *μm*, total volume 0.261 *μm*^3^ and surface area 1.571 *μm*^2^. The total tetrahedron number is 1394, and the mean tetrahedron edge length is 124 nm, with maximum and minimum edge lengths of 161 nm and 84 nm, respectively. A total of 300 vesicles with a diameter of 40 nm are present. As initial condition a portion of vesicles are docked in active zones (AZs), the rest of the vesicles are randomly distributed in the mesh and diffuse or are actively transported by actin (tether) to dock into the active zones. Each docked triangle in the active zone has one tether at the normal direction 100 nm long with a transport velocity of 200 µm/min (Gallimore et al., 2025; Hepburn et al., 2024). Based on previous observations (Lu & Trussell, 2000; Otsu et al., 2004), we assume that SR and AR are derived from the same pool of release-ready vesicles.

We assume the same number of SNARE_CXN_syt7 complexes initially. Later the number of syt7 complexes is increased, conforming to experimental observations in hippocampus (Bacaj et al., 2013; Sugita et al., 2001; Voleti et al., 2017). In general, the relative number of syt1 and syt7 complexes might differ between neurons and synapses, and this will affect the release patterns of SR and AR (Turecek & Regehr, 2019).

Calcium ion influx is mediated by voltage-dependent CaP-type channels described as a GHK current, with variable numbers of channels on each docking triangle. In many simulations we use 1.5 calcium channels per docking triangle, which means that randomly either 1 or 2 channels are placed on each docking triangle. There are 4 AZs with a total of 114 docking triangles, each AZ has an area of 0.11 *μm*^2^. The calcum pumps have a density of *ρ_pump_* = 6.022 *μm*^−2^ on the membrane. We have not included the mechanism of Calcium Induced Calcium Release (CICR) from endoplasmic reticulum. The endogeneous buffers include PV, CaM, CB with high (CBhi) and low affinity (CBlo) and CRTT with concentrations as in (Gallimore et al., 2025). Note that the CaM has a high concentration of 60 µM (Gallimore et al., 2025; Wilhelm et al., 2014).

Two stimulation paradigms are used: a clamped calcium concentration at different concentrations or 10 depolarization pulses at frequencies of 10, 20, 50, or 100Hz. Membrane potential is clamped to 40 mV for 1 ms for each depolarization pulse and clamped to the resting potential of -61 mV when there is no pulse.

## 3. Results

### 3.1. Syt7 has a higher intrinsic affinity than syt1 and a slower timescale

To characterize the intrinsic affinity of syt7, kinetic parameters of calcium binding and unbinding process were optimized (Table 1). As can be seen in Fig. 2C, syt7 exhibits an intrinsic affinity of about 10 µM (Methods 2.1.3), whereas syt1 has a much higher intrinsic affinity of about 61µM (Hui et al., 2005). CaM (Gallimore et al., 2025) has an intrinsic affinity similar to that of syt7. The apparent affinity based on the intrinsic affinity was fit to experimental observations (Hui et al., 2005; Tran et al., 2019) shown by the colored dots in Fig. 2A, 2B. Note that apparent affinity is higher than the intrinsic one for both syts, but the difference between the two syts persists.

The speed of calcium ion binding for syt7, syt1 and CaM is compared in Fig. 2D, using the effective calcium binding timescales (Methods 2.1.4). As shown, syt1 binds calcium about 1-3 orders of magnitude faster than syt7 at various calcium concentrations, making syt7 a much slower sensor. Syt1 is also much faster than CaM. Syt7 becomes faster than CaM at calcium concentrations higher than 1 µM. In addition to the calcium binding kinetics, the experimentally observed exocytosis rate of syt1 is faster than syt7 (Davis et al., 1999; Hui et al., 2005; Tran et al., 2019) resulting in a much higher release rate parameter *γ*_2_ for syt1 (Methods).

Next, we use the models of syt1 and syt7 to explore release patterns in a model of a synaptic bouton (Fig.1F) in STEPS.

### 3.2. Vesicle release switches from syt7 to syt1 dominance at high calcium

Using calcium clamped simulations lasting 0.5 s, the pseudo apparent affinity of syt1 is around 6.0 µM (Fig. 3A blue, syt7KO) whereas that of syt7 is around 3.0 µM (Fig. 3A orange, syt1KO). For syt7, our results are in the range of experimental results (Fukuda et al., 2004; Sugita et al., 2002). With coexistent syt1 and syt7 in equal amounts (4 each on a vesicle), our results show a critical calcium concentration of about 9 µM to switch from syt7-triggered to syt1-triggered release (Fig. 3B). This critical calcium level is independent on the total number of vesicles (tested by doubling the number of vesicles), indicating that the intrinsic properties of syt1 and syt7 are the fundamental factors that determine the outcome of their competition in triggering vesicle release. Detailed examples for vesicle release and events at each calcium clamped value are shown in Fig. 3C, D. For calcium concentrations lower than 9 µM, syt7 triggered more vesicle releases; at Ca = 9 µM , syt1 and syt7 released similar number of vesicles, but syt1 is faster than syt7; for Ca = 15 µM, syt1 triggered many more releases with faster speed than syt7.

**Figure 3.**
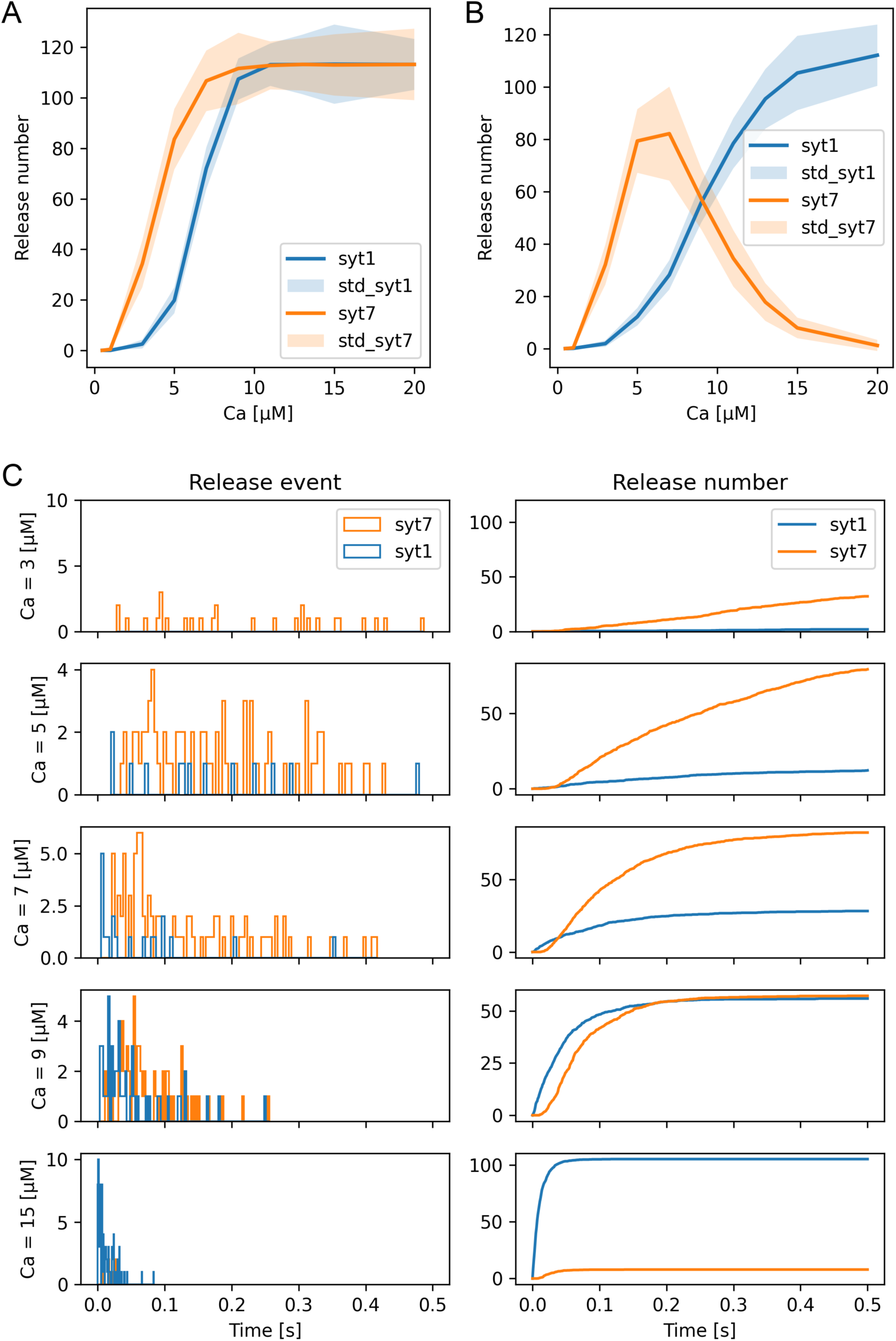
(A, B) Dependence of pseudo apparent affinity of syt1 and syt7 to calcium concentration clamped simulations of release from a synaptic bouton for syt1 or syt7 Knock Out (KO) (A) or syt1 and syt7 coexisting condition (B). (C) Examples of dynamics of vesicle release events and accumulated number of releases for indicated concentration of calcium clamp. Standard parameters, mean of 10 iterations.

We observe that syt7 triggers less vesicle release in the syt1 coexisting condition (Fig. 3B) than syt1KO (Fig. 3A) for calcium levels higher than 10 µM, which is consistent with experimental observations that syt1KO facilitates syt7’s release (Bacaj et al., 2013; Maximov & Südhof, 2005). Due to the faster speed of exocytosis by syt1 and saturation of calcium binding for syt1, syt7-triggered vesicle release drops dramatically at calcium concentrations higher than 9 µM.

In the following sections, we extend these observations to release caused by voltage pulses of different frequencies.

### 3.3. Competition between syt1 and syt7 triggers SR and AR by voltage pulses

In Fig. 4 we show an example simulation of the competition between syt7-triggered AR and syt1-triggered SR caused by 10 pulses at 50 Hz. The pattern of syt1-triggered SR is tightly coupled to the voltage pulses in time (within 3 ms of each spike) (Bacaj et al., 2013; Kaeser & Regehr, 2014; Südhof, 2013) ,while syt7-triggered vesicle release occurs during the stimulus train (following 3 ms after the spike), and after the termination of stimuli at 0.2s, i.e. AR (Fig. 4D). We show in Fig.4-figure supplement 1 that syt1-triggered release is mainly SR and syt7-triggered release is mainly AR. For syt1 most of the vesicles are released as 2 SNARE_CXN_syt1, much less as 3 SNARE_CXN_syt1 complex and almost none as 4 SNARE_CXN_syt1 (Fig. 4B), indicating non-saturation for calcium binding. Conversely, syt7 release has the highest contribution from 3 SNARE_CXN_syt7 complex with also a sizable 4SNARE_CXN_syt7 component (Fig. 4C), indicating more effective calcium binding due to its higher intrinsic affinity. As expected, the kinetics of syt1 C2A and C2B domains in binding and unbinding calcium (Fig. 4F) are much faster than those of syt7 (Fig. 4G), such that after each spike, Ca bound syt1 C2A domains drop to zero fast, providing no chance for AR. Syt7 binds and unbinds calcium much slower, allowing for AR after terminating stimulation. Finally, PV (Fig. 4E) and CaM (Fig. 4F) buffer large amounts of calcium, PV_2Ca reaches 13 µM and maximum of CaM[C2,N2] is about 30 µM. Since CaM unbinds calcium much faster than PV, CaM releases large amounts of calcium to the cytosol after stimulation ends, causing a slower decay of the calcium concentration, often referred to as “residual calcium” (Chamberland et al., 2020; Timofeeva & Volynski, 2015).

**Figure 4.**
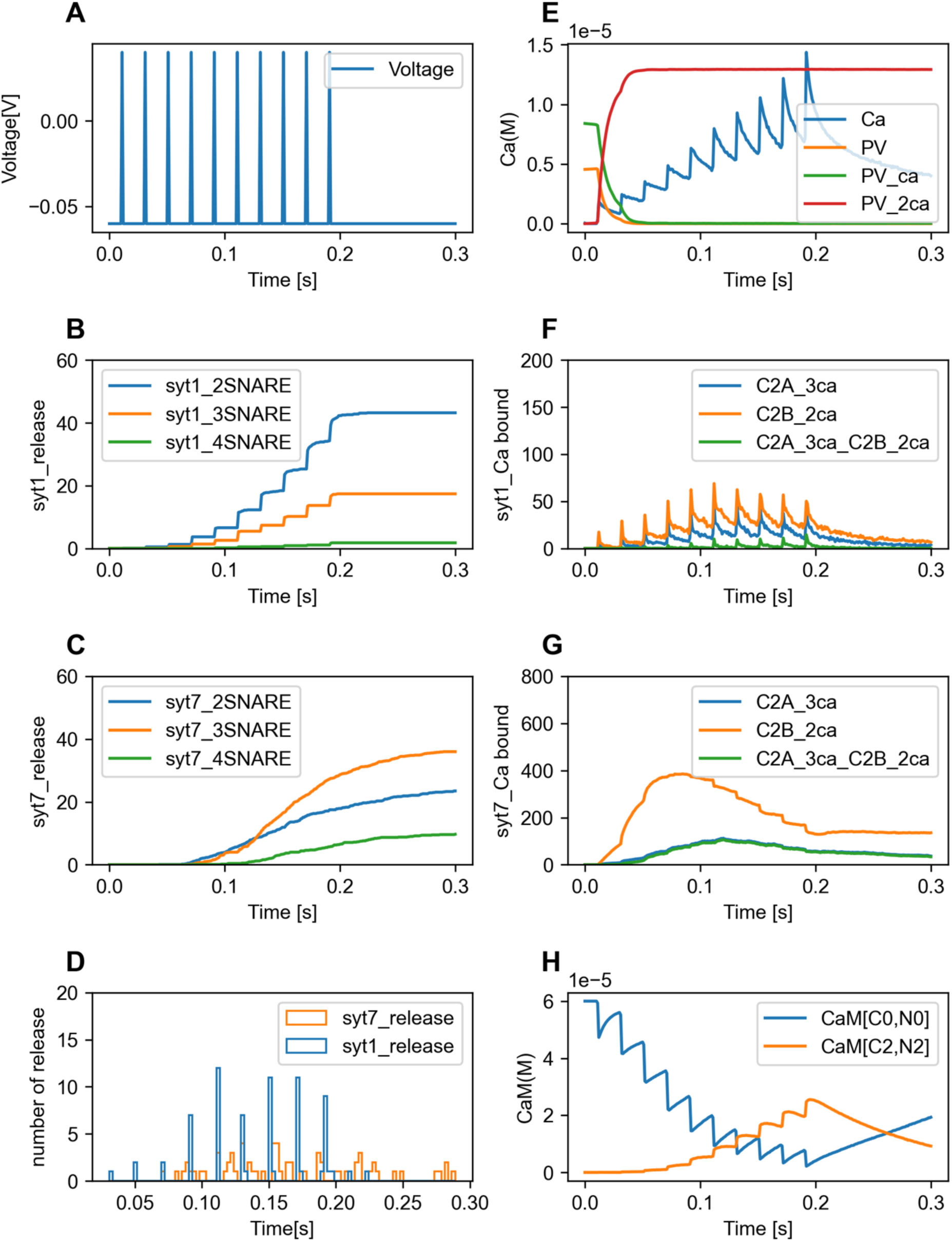
Example of syt1-triggered SR and syt7-triggered AR by 10 pulse stimuli at 50 Hz. (A) voltage dynamics. (B,C) vesicle release by 2,3,4 SNARE complex for syt1 (B) and syt7 (C). (D) release events caused by syt1 and syt7. (E) dynamics of free calcium and buffer PV (F,G) kinetics of calcium binding by C2A and C2B domains for syt1 (F) and syt7 (G). (H) kinetics of calcium binding by CaM: [C0,N0] no calcium bound, [C2,N2] fully bound. Parameters: syt1 = 4, syt7 = 4, CaM = 60 µM, 1.5 Cachan/dock_tri, Ca_ocon = 2 mM, mean of 10 iterations.

Next, we investigate how frequency of stimulation affects release properties.

### 3.4. Syt1KO facilitates syt7-triggered release at specific conditions

Fig. 5A shows the response of the standard model to 10 voltage pulses at 100 Hz. At this rate, with coexistent syt1 and syt7, SR by syt1 dominates vesicle release (Fig. 5A2, A4) and syt7 causes release of only 28 vesicles. Conversely, in a syt1KO model syt7-triggered vesicle is facilitated (Fig 5B), causing AR of 100 vesicles (Fig. 5B2). This can be explained by the observation that the C2A and C2B domains of syt7 in the syt1KO model have max 160 fully calcium bound state (Fig. 5B3), whereas in the coexistent condition they have only max 100 copies (Fig. 5A3). Notice that the free calcium concentration dynamics look almost identical for both simulations (Fig. 5A1 and 5B1), indicating that the free calcium dynamics do not predict the release pattern or which sensors are involved, which will be illustrated in more detail in the following sections.

**Figure 5.**
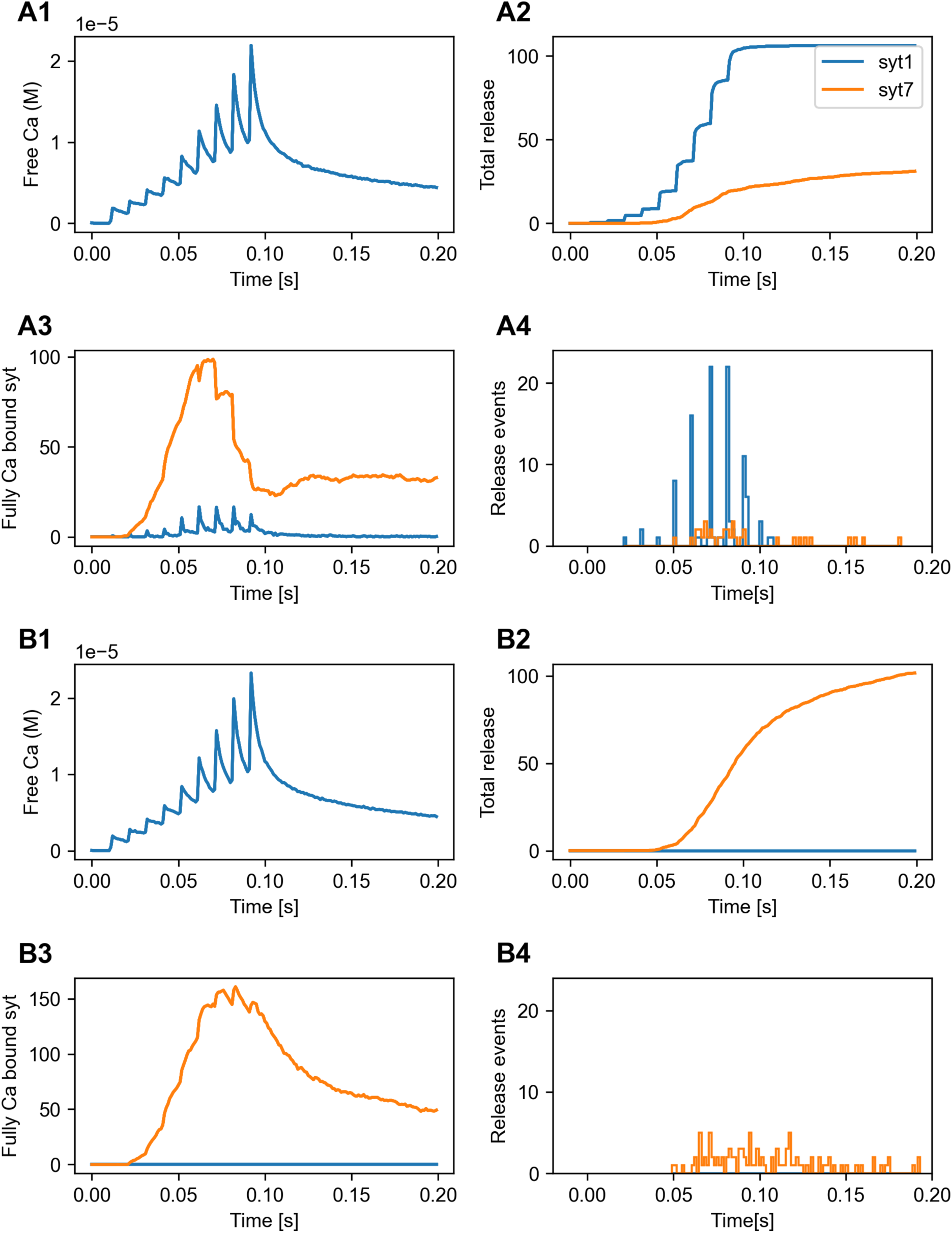
Example simulation showing that syt1KO facilitates syt7-triggered release. (A1-A4): coexistence of syt1 and syt7; (B1-B4): syt1KO condition. (A1, B1) free calcium dynamics caused by 10 voltage pulses at 100 Hz. (A2,B2) vesicle releasing dynamics. (A3, B3) dynamics of fully calcium bound syt numbers. (A4, B4) vesicle release events triggered by syt1 and syt7. Parameters: syt1 = 4, syt7 = 4, CaM = 60 µM, 1.5 Cachan/dock_tri, Ca_ocon = 2 mM, mean of 10 iterations.

In Fig. 6 we compare the responses to different stimulation frequencies for the complete model to both KO conditions for low calcium influx (Fig. 6A: 1 calcium channel per docking triangle) or high calcium influx (Fig. 6B: 1.5 channels per docking triangle). For the low calcium condition, syt7 in the syt1KO condition (A2) triggers similar vesicle releases to the coexisting condition (A1) for all frequencies except for 100Hz, because the calcium level is not high enough for syt1 to become competitive to syt7. Conversely, in the syt7KO (A3) condition, the lack of competition allows syt1 to trigger higher vesicle releases for high stimulation frequencies. The KO conditions have a much stronger effect in the high calcium influx condition (B2, B3). Now the lack of competition allows the remaining syt to cause significantly higher release at all frequencies except 10 Hz.

**Figure 6.**
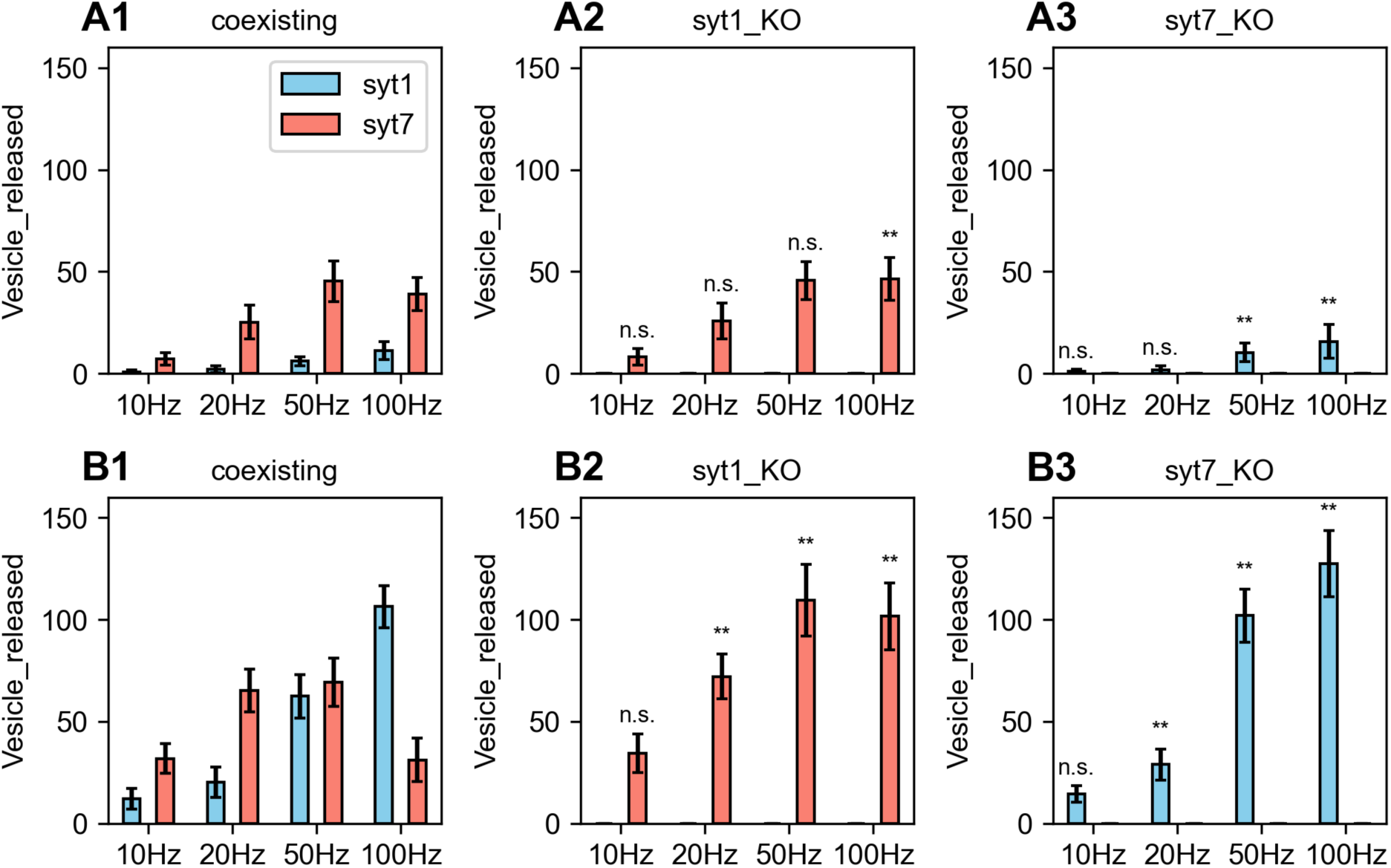
Syt1KO facilitates syt7-triggered AR only under specific conditions. Both syt7KO and syt1KO can show frequency dependent increases. (A1, A2, A3) 1 Cachan/docking_triangle (B1, B2, B3) 1.5 Cachan/docking_triangle. (A1, B1) coexisting syt1 = 4 and syt7 = 4. (A2, B2) syt1KO. (A3, B3) syt7KO. Other parameters: CaM = 60 µM, Ca_ocon = 2 mM, mean of 10 iterations. Statistics: comparison of KO to coexisting condition, ** p<0.05, n.s not significant (p>0.05).

It is important to point out that the cytosolic free calcium level does not predict the release kinetics, or even which sensor is involved. Fig. 6-figure supplement 1 shows the response to 50 Hz in Fig. 6B in more detail. The cytosolic calcium concentration (Fig. 6-figure supplement 1 A2, B2, C2) and buffering (A4, B4, C4) dynamics are indistinguishable for different sensor conditions (A: coexist; B: syt1KO; C: syt7KO), but the release patterns are totally distinct (A3, B3, C3).

### 3.5 Effect of calcium channel distribution on local concentrations and sensor competition

The comparison between Fig. 6A and 6B indicates that local calcium concentrations may play a key role in determining release patterns. To further explore these effects, we compare the standard model with uniform distribution of calcium channels to an extremely clustered case with all calcium channels in a single docking triangle (Fig. 7). For a low number of 114 calcium channels (A1, A2), the uniform distribution condition (A1) shows more syt7-triggered AR, and the syt1-triggered release is quite small. Conversely, for the clustered calcium channels (A2), syt1-triggered release increases dramatically for all frequencies, while syt7-triggered release is a bit smaller. This indicates that the clustered distribution of calcium channels favors syt1-triggered SR. One example of detailed comparisons between uniform and clustered distribution of calcium channels in global and local calcium dynamics as well as release kinetics can be seen in Fig. 7-figure supplement 1, where the global free calcium dynamics is almost the same, but uniform distribution mainly exhibits syt7-triggered AR, while the clustered distribution shows both syt1 triggered release and syt7-triggered release with almost the same magnitude.

**Figure 7.**
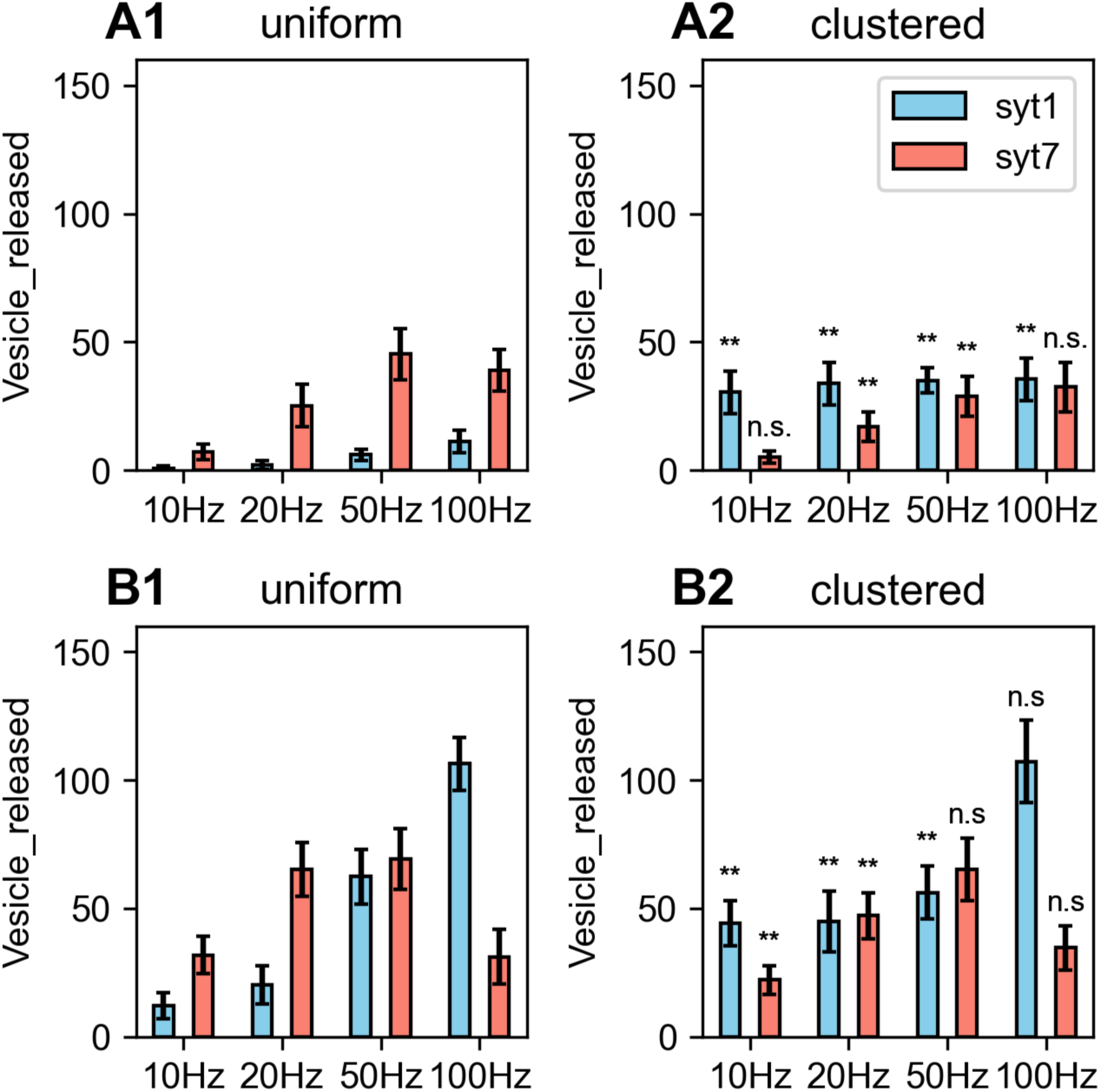
Calcium channel distribution controls release patterns: uniform versus clustered. (a1, a2) total number of calcium channel is 114, with (A1) uniformly 1 calcium channel in each of 114 docking triangles and (A2) all 114 calcium channels in a single docking triangle. (B1-B2) same for 171 calcium channels, with (B1) uniformly 1.5 calcium channels per docking triangle and (B2): all 171 calcium channels in a single docking triangle. Parameters: syt1 = 4, syt7 = 4, CaM = 60 µM, Ca_ocon = 2 mM, mean of 10 iterations. Statistics: comparison of clustered to uniform condition, ** p<0.05, n.s not significant (p>0.05).

For a higher number of calcium channels of 171, the differences between the uniform and clustered distribution (Fig. 7B) become smaller and are only significant for lower stimulation frequencies. This indicates that a clustered channel distribution can facilitate syt1-triggered release, especially at low channel numbers and at low frequencies.

A more realistic clustering assumption is to have multiple clusters. This is investigated in Fig. 7-figure supplement 2, where the total number of calcium number is kept fixed at 114 and the number of channel clusters decreases with cluster size. Similar to Fig. 7 we observe more facilitation of syt1-triggered release at a low frequency (Fig. 7-figure supplement 2 A,B,C), but this effect saturates at 16 calcium channels per docking triangle (D,E,F).

The effect of calcium channel distribution on the release patterns is affected by CaM buffering large amounts of free calcium, so that the syt sensors have to compete with it. For example, for 114 channel all in a single docking triangle clustered condition, the release is dominated by syt1-triggered release with CaM = 0 µM (Fig. 8 A1) for all frequencies, but with CaM = 60 µM (Fig. 8 A2) syt1-triggered release drops dramatically. Therefore, CaM competes with syt1 for calcium binding, whereas syt7 triggered release is much less affected by CaM and even increases at 100Hz stimuli due to the CaM modifying the competition between syt1 and syt7. These results can be explained by the fact that syt7 and CaM have similar intrinsic affinities with a KD about 10 µM whereas syt1 has much lower affinity with a KD about 60 µM.

**Figure 8.**
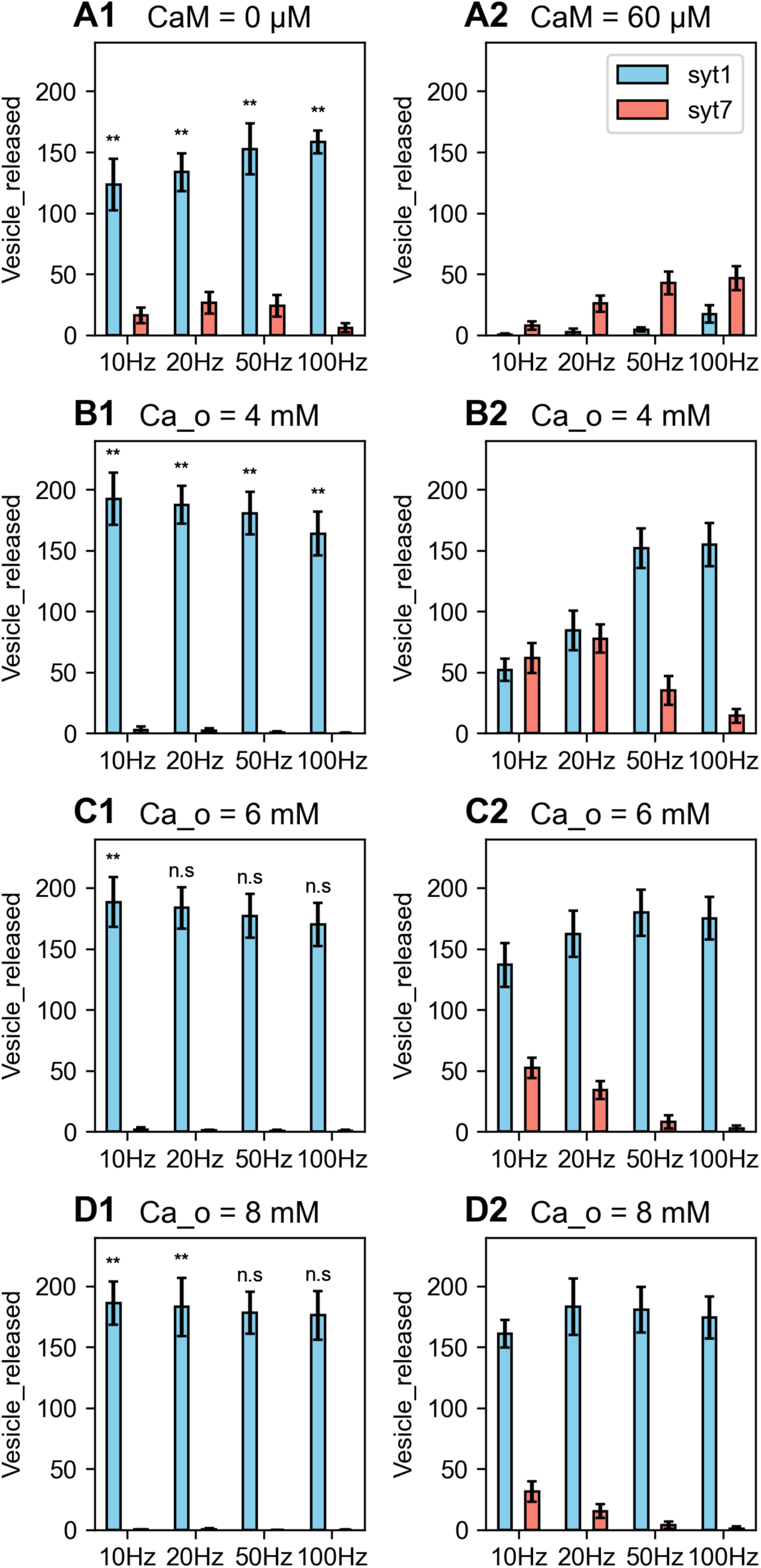
Syt1 and syt7-triggered release depend on presence of CaM and on calcium external concentration, Ca_ocon. (A, B, C, D) Ca_ocon = 2 mM, 4 mM, 6 mM, 8 mM. (A1, B1, C1, D1) CaM = 0 µM; (A2, B2, C2, D2) CaM = 60 µM. Parameters: syt1 = 4, syt7 = 4, 1 Cachan/dock_tri, mean of 10 iterations. Statistics: black ** (p<0.05) and ‘n.s.’ in (A1,B1,C1,D1) shows the pair-wise comparison of vesicle release triggered by syt1 between CaM = 0 µM (left column) and CaM = 60 µM (right column).

The effect of CaM depends on the external calcium concentration. When it is increased to 6 mM or 8 mM as shown in Fig. 8 C, D, the cytosolic calcium concentration also increases (see Fig. 8-figure supplement 1), thus CaM buffering will be saturated, therefore, syt1-trigged release becomes less affected by the existence of CaM = 60 µM (Fig. 8 C2, D2).

In Fig. 9 the buffering effect of CaM on the time-dependent spatial distribution of free calcium is shown. The free calcium level drops dramatically by buffering with CaM = 60 µM (A2, B2) at t = 0.012 second (green dots) compared to CaM = 0 µM (A1, B1). We can see that the spatial constant of this buffering effect is about 300 nm from the point of calcium entry. The theoretical predictions for CaM are *τ* = 0.216 *ms* (*eq*. 1) and *λ* = 0.22 *μm* (*eq*. 2) and for faster buffer PV *τ* = 0.623 *ms* and *λ* = 0.37 *μm*. Both of these values are within the range of the effects observed in Fig. 9.

**Figure 9.**
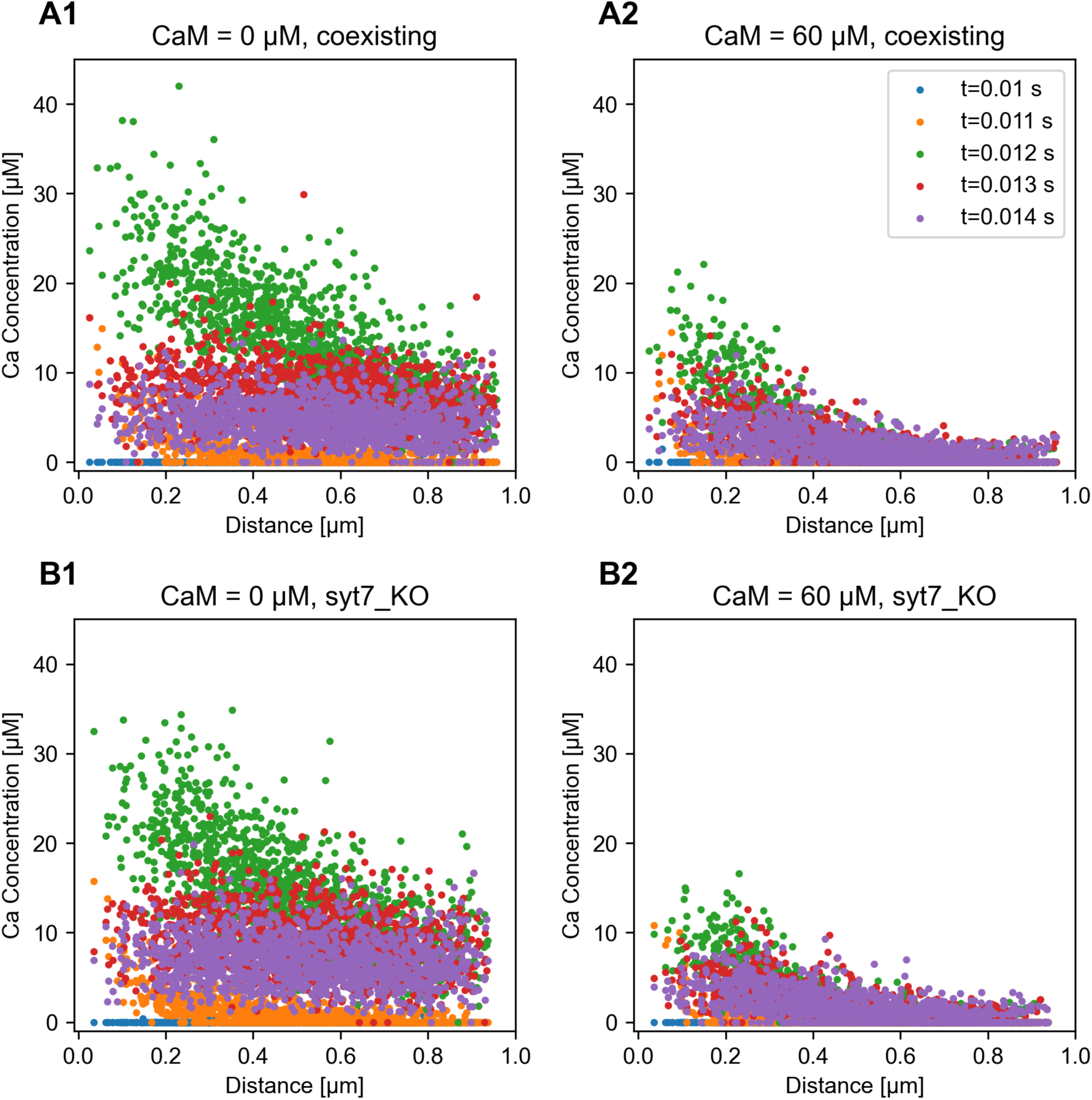
CaM buffers calcium in a sub-millisecond range with a strong effect at a spatial distance of 200 nm or more. (A1,A2) coexisting condition. (B1,B2) syt7_KO. (A1, B1) CaM = 0 µM; (A2, B2) CaM = 60 µM. All 171 calcium channels are put in a single docking triangle. Blue dots: at rest (voltage = - 61 mV, t = 0.010 s). orange dots: 1 ms after of voltage jump to 40 mV (t = 0.011 s) , green dots: immediately after voltage is clamped back to rest at -61 mV (t = 0.012 s), red dots: 1 ms later at -61 mV (t = 0.013 s), purple dots: t = 0.014 s. Data points are calcium concentrations for each tetrahedron at the indicated distances from the location of calcium influx, and the data are the average value of 10 iterations. Parameters: Ca_ocon = 2 mM, 100Hz stimuli with 10 pulses.

Therefore, our simulation results of Fig. 7 can be explained. The clustering of calcium channels causes a higher magnitude of calcium influx and will facilitate syt1-triggered release if the clusters are closer to the vesicle than the buffering spatial constant *λ*.

Otherwise, with a uniform distribution of calcium channels which has lower calcium influx, CaM will quickly buffer free calcium due to its high concentrations and prevent syt1 from binding enough calcium to trigger SR, due to higher KD of syt1 than CaM, especially with low frequency stimuli and low calcium channel numbers. In contrast, syt7 can still trigger AR, because its KD is comparable to that of CaM.

### 3.6 Effect of relative number of syt7 and syt1 on releasing patterns

For all previous simulations, we had syt1 = syt7 = 4 for equal competition. But, based on experimental observations, syt7 gene expression is almost triple that of syt1 in hippocampal neurons (Bacaj et al., 2013). Therefore, we investigate the effect of simulations with syt7 = 12 and syt1 = 4 in this section.

In Fig. 10, the tripling of the syt7 amount with clustered calcium channels causes much more release for all frequencies (Fig. 10). For small calcium channel cluster sizes this enhances the frequency dependence of syt7 release (A2,B2), but the effect saturates at higher number of calcium channels (C2).

**Figure 10.**
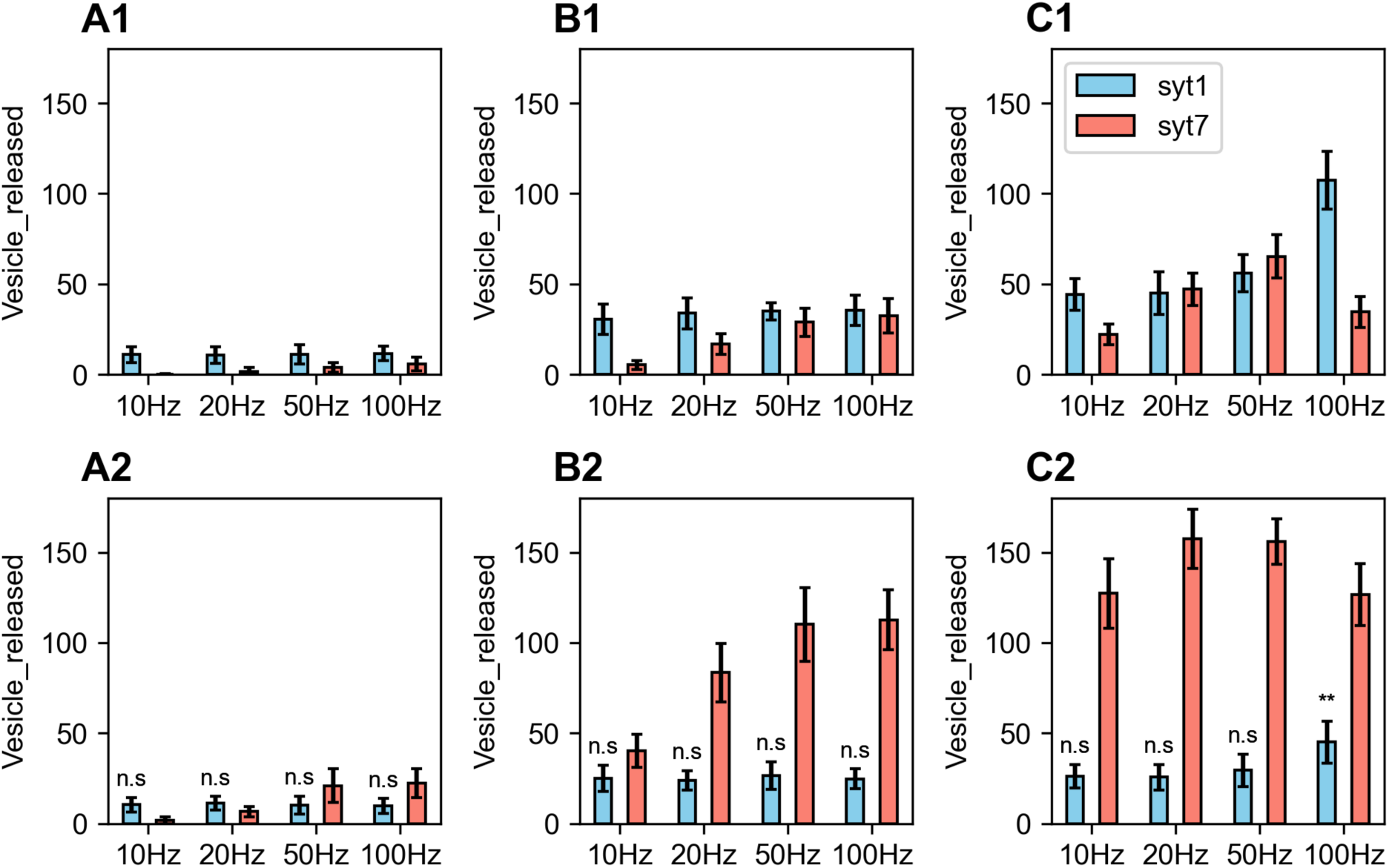
Frequency encoding for syt7 is more pronounced when number of syt7 is tripled, and syt1-triggered frequency independent release. (A1-C1) syt7 = 4, syt1 = 4. (A2-C2) syt7 = 12, syt1 = 4. Calcium channel number in a single docking triangle = 57, 114, 171 for (A, B, C) respectively. Other parameters: CaM = 60 µM, Ca_ocon = 2 mM, mean of 10 iterations. Statistics: syt1-triggered release show no significant difference by different frequencies of stimuli within each panel of A2,B2,C2. ** p<0.05, n.s not significant (p>0.05). In C2, 100 Hz show significant difference from 10Hz, 20Hz, 50Hz, respectively.

Conversely, syt1 -triggered release is frequency independent for almost all cluster sizes at both syt7 = 4 and syt7 = 12 conditions (Fig. 10). This indicates that the calcium concentrations sensed by syt1 decay at a timescale faster than the inter spike intervals. This is due to the high KD of syt1 and large amount of CaM with a fast-buffering effect that prevents syt1 of binding more calcium ions to trigger extra release at higher frequency stimuli. Whereas for syt7, with a similar KD as CaM, it has a higher chance to bind calcium with higher frequency stimuli, especially if its quantity is tripled.

Fig. 10-figure supplement 1 shows more detailed data for the simulations in Fig.10 B2. Note that the global calcium concentration increases with high frequencies without affecting the syt1-triggered release, while syt7-trigggered release increases with frequency. Fig. 10- figure supplement 2 plots, similarly to Fig. 9, the effect of cluster size on calcium spatial and temporal dynamics.

We conclude that the competition between syt1 and syt7 in binding calcium is the key mechanism to explain differential release patterns, and their competition can be regulated by CaM through its buffering effect. The steady-state partitioning of calcium (eq. 3) between each syt sensors and CaM and their relationship with free calcium concentration is shown in Fig. 10-figure supplement 3, which illustrates the importance of the dissociation constant and sensor concentration in partitioning calcium ions (see SI Section 4), leading to different patterns of vesicle release.

## 4. Discussion

We developed a quantitative model of synaptotagmin-7 calcium-binding kinetics and integrated it with synaptotagmin-1 in a half-hemisphere mesh to simulate vesicle release from a presynaptic bouton. To capture syt7 calcium binding properties, we treated the C2A and C2B domains as separate binding modules with distinct kinetics (Sugita et al., 2001, 2002; Zhou et al., 2015, 2017). Rate constants were constrained by intrinsic and apparent calcium affinities based on experimental data (Tran et al., 2019; Voleti et al., 2017), and we derived quantitative expression of intrinsic affinity as a function of the ratio of *k*_off_/*k*_on_ and apparent affinity by incorporating lipid interactions (see SI Section 3), to better characterize the sensor property. Within this framework, the timing of calcium binding—quantified by an effective timescale *τ* for reaching full occupancy—as well as the releasing rates emerge as key determinants of vesicle release kinetics.

Our simulations reveal a critical calcium threshold (∼10 μM) above which syt1 tends to outcompete syt7 in driving vesicle release. Consistent with experimental observations (Bacaj et al.,2013) syt1 knockout enhances syt7-mediated release under specific calcium-clamped and pulse conditions, while having limited effects in other regimes. Under clustered calcium channel configurations with elevated syt7 abundance, syt7-mediated release increases with stimulation frequency, whereas syt1-mediated release remains relatively frequency-independent, which provides a potential intepretation for spike-counting mechanism in MFB (Chamberland et al., 2018).

Mechanistically, these behaviors arise from the interplay between calcium-binding kinetics and downstream release rates. Lipid association and dissociation kinetics are faster for syt1 than for syt7 (Hui et al., 2005), and are represented here through effective release rates. Under saturating calcium, syt1 triggers vesicle fusion more rapidly (*γ*_2_ = 3000 s^−1^v.s. 10 s^−1^for syt7), leading to syt1-dominated release. In contrast, under sub-saturating conditions, calcium-binding kinetics become rate-limiting, and sensor-specific affinities govern release dynamics. This transition between regimes is captured by two characteristic timescales in the underlying reaction scheme (see SI Section 2).

In addition, the endogenous buffer CaM plays a central role by competing with synaptotagmin sensors for calcium binding. Because CaM has a higher affinity than syt1 but a comparable affinity to syt7, it preferentially suppresses syt1-mediated release with a more limited effect on syt7. Importantly, our results show that free calcium concentrations alone do not predict release mode; rather, release dynamics are determined by how calcium is partitioned among sensors and buffers. This partitioning depends on both dissociation constants and relative abundance (see SI Section 4), providing a mechanistic basis for diverse synaptic release patterns.

Our model uses a simplified half-hemisphere geometry to approximate a presynaptic bouton. Although this abstraction does not fully capture native morphology, it reproduces key features, including a readily releasable pool of several hundred vesicles and a realistic active zone area (∼0.11 µm²). Vesicles are assumed to be initially docked, with an additional fraction undergoing docking when the sites become free after release. While this allows us to isolate calcium-dependent release mechanisms, it does not account for vesicle recycling or endocytosis, which are likely to influence release dynamics during sustained activity (Gallimore et al., 2025). Extending the model to include these processes will be important for understanding their roles in short-term and longer-term synaptic plasticity.

We also simplify the molecular reactions leading to fusion by assuming that vesicles are pre-associated with SNARE–synaptotagmin–complexin complexes, thereby bypassing explicit modeling of SNARE binding. However, emerging evidence suggests that syt1 and syt7 may compete for SNARE binding and form distinct fusion-competent complexes (Bose et al., 2024). Given their different subcellular localizations, such interactions could introduce additional kinetic constraints and regulatory pathways. Incorporating these reactions into a more complete network model may reveal additional modes of release dynamics and further diversify synaptic responses.

Finally, complexity arises from the diverse roles of syt7 in regulating vesicle release (Fukuda et al., 2004; Huson & Regehr, 2020; Liu et al., 2014; MacDougall et al., 2018; Wang et al., 2005). Syt7 is required for short-term facilitation (Jackman et al., 2016) and mediates both facilitation and asynchronous release at granule cell synapses (Turecek & Regehr, 2018). Moreover, recent studies showed that introduction of syt7 promotes facilitation at climbing fiber–Purkinje cell synapses (Weyrer et al., 2021). In addition, asynchronous release may involve other calcium sensors beyond synaptotagmins (Wu et al., 2024; Yao et al., 2011). While our study emphasizes competition between syt1 and Sst7, elucidating how competition and cooperation are coordinated represents an important direction for future work, and may provide further insight into how synapses flexibly tune transmission and plasticity.

## Supplementary Information

### 1. The full solution for the 2-state model *A* ↔ *B*

We assume the solutions are written in the following form:

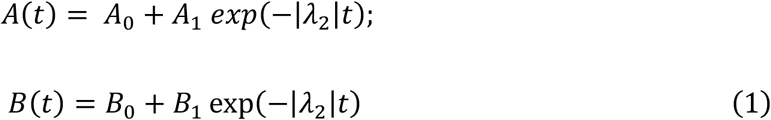

By *A*(*t* = 0) = 1, *A*(*t* → ∞) = *c*_A_(*equilibrium concentration*), then we can get: *A*_0_ + *A*_1_ = 1; *A*_0_ = *c*_A_.

therefore, we can get:

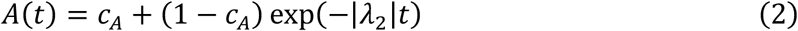

Similarly, *B*(*t* = 0) = 0, *B*(*t* → ∞) = *c_B_*(*equilibrium concentration*)

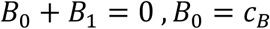

we can see that

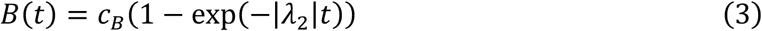

*w*ℎ*ere* 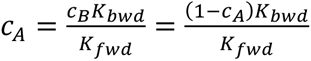, is the equilibrium solution, thus

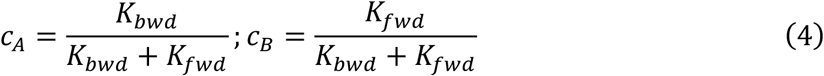

## 2. Exact solution for a theoretical 3-state model

The model of A binds calcium and goes to B state, where it triggers release to C state with *γ*_2_can be expressed as:

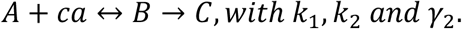

The model can be mapped to syt model as: syt effectively calcium binding rate *K_f_*_w*d*_ and effective unbinding rate *K_b_*_w*d*_, by this reaction in linear releasing probability:

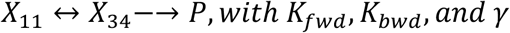

So, the dynamic equation can be written as:

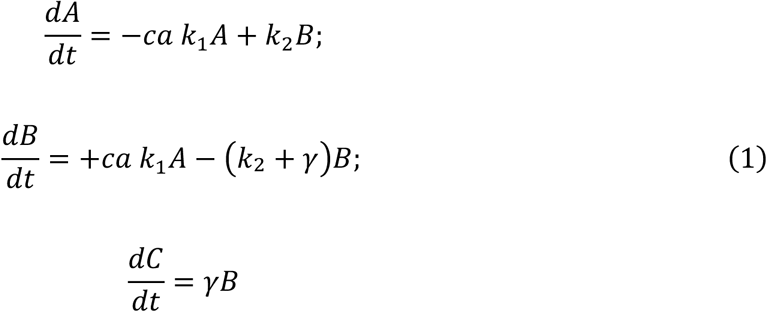

Correspondingly, the eigen value can be calculated by:

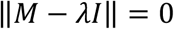

And we obtain 3 eigen values:

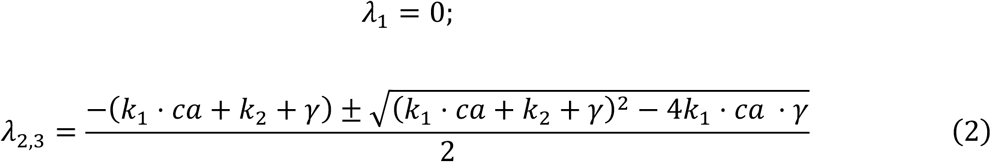

We can now write the solution of *A*, *B*, *C* by:

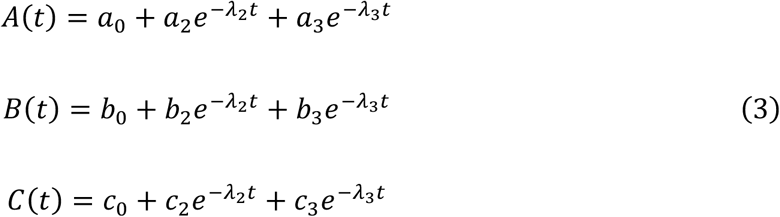

By the initial condition,*t* = 0, *A* = 1, *B* = 0, *C* = 0, we have:

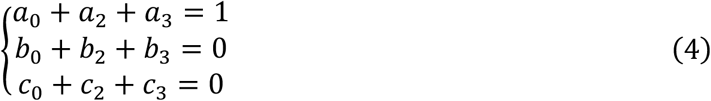

By infinity condition,*t* → ∞, *A* = 0, *B* = 0, *C* = 1, we have:

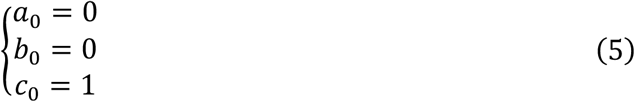

By the conservation law, *A* + *B* + *C* = 1, we now have:

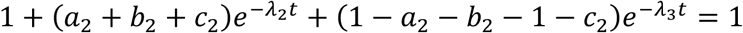

Which is

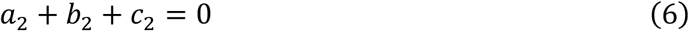

So far, we have 9 unknowns and 7 constraints, still we need another constraint, that is:

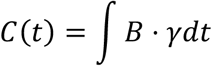

Which yields:

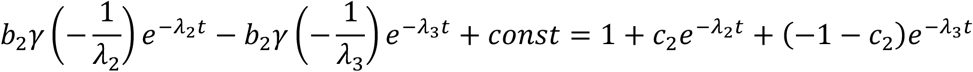

By comparing the coefficients of exponential term, we can get two relations:

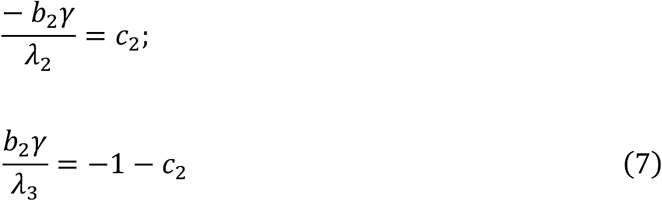

So eventually we have 9 constraints of eq. 1 to 4, and the coefficients can all be expressed by the gamma and lambdas:

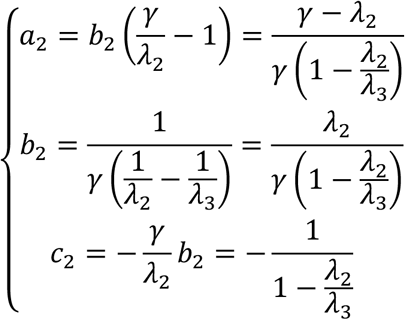

Now, we have the full expression of the 3-state model.

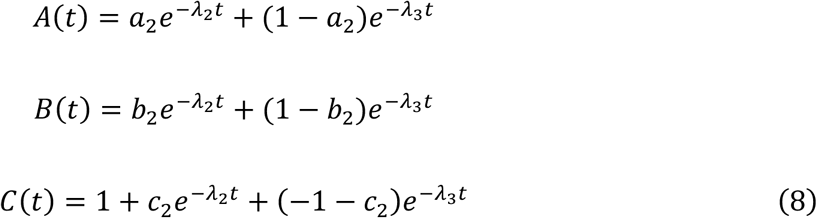

There are two points need to address:

1. only with linear releasing condition, can we get the exact solution. If using 2 SNAREs with *γ*_2_ × *S_fCa_*^2^ nonlinear terms, there will be no eigenvalues computed from above calculations.
2. By putting *k*_1_*ca*−→ *K_f_*_w*d*_, *k*_2_−→ *K_b_*_w*d*_, *γ*−→ *γ*_2_, we can theoretically analyze the behavior of calcium binding and releasing kinetics in principle. Nevertheless, this is still an approximation, since the effective rates *K_f_*_w*d*,_ *K_b_*_w*d*_ speed up the original calcium binding reactions.

## 3. The apparent affinity for syt1 and syt7

By adding the lipid binding kinetics, the equilibrium state will be written as a function of lipid concentration *lip_c_*, and lipid binding and unbinding rates of *lip_on_* and *lip_off_*:

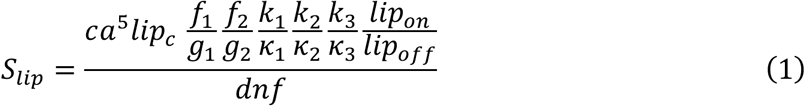

Where the denominator factor *dnf* is:

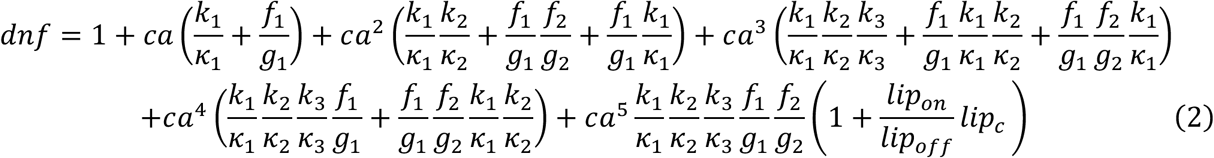

And *S_lip_* is a function of calcium concentration. Given *lip_on_* , *lip_off_*, *lip_c_* we can scan the calcium concentrations and obtain a sigmoid curve for *S_lip_* (which is named “calcium dependent lipid binding” in the literature), which can be used to determine the apparent affinity of each sensor (eq. 1). This is our theoretical derivations based on detailed balance solution.

## 4. Partition of calcium by each sensor and buffer at equilibrium state

Here we analytically calculate the partition of calcium for syt1 (A) and syt7 (B) and CaM (C) at equilibrium state at calcium clamped conditions.

We assume initial calcium concentration is [*Ca*^0^], which is conserved.

Initial concentration of A, B, C is [*A*_0_], [*B*_0_], [*C*_0_], the chemical reactions can be simply shown here:

A + Ca ↔ [CaA]

B + Ca ↔ [CaB]

C + Ca ↔ [CaC]

And the dynamics equation of the kinetics is shown as:

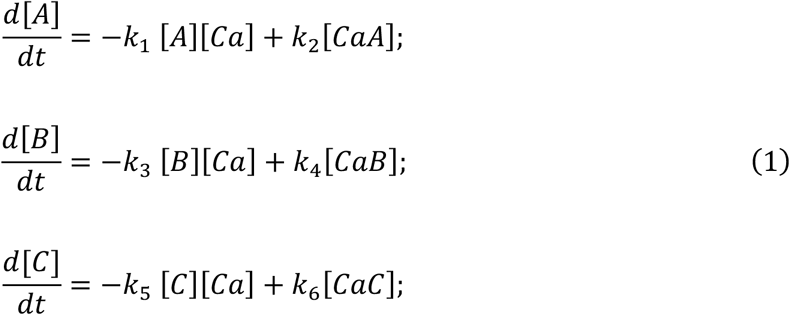

By the equilibrium condition, we have the detailed balance solution:

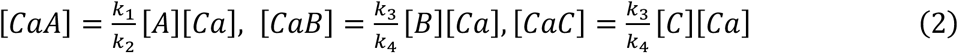

By the conservation of calcium, we can write the following equation:

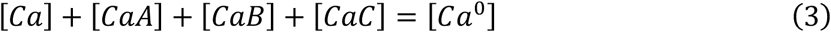

By using the conservation of species, A , B, C we can also obtain the relation:

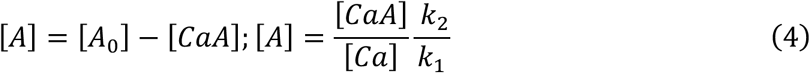

From eq. 4 we obtain:

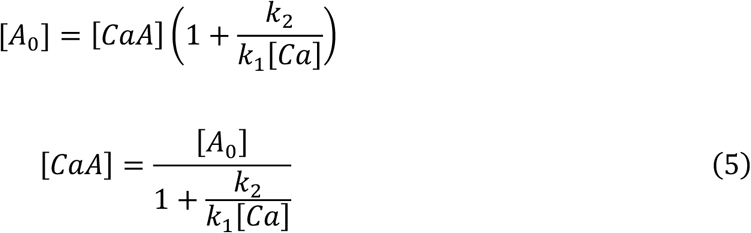

So, we can write similar expression for the [CaB] and [CaC] and put them into eq. 3 to

obtain:

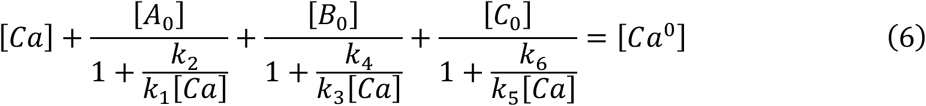

So, eq. 6 illustrates that at equilibrium state, the free calcium [Ca] and the calcium occupied by each species of A, B, C can be computed. We call this the partitioning of calcium.

We can write eq. 6 in a new form:

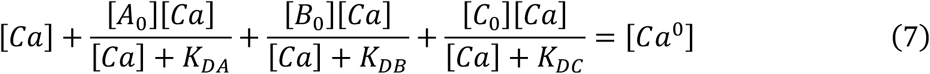

where, the term of 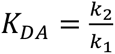 is the dissociation constant of species A by definition. Same for *K_DB_*, *K_DC_*.

From eq. 7, we see that if A has a very high *K_D_*_A_ ≫ [*Ca*] it will not capture a lot of calcium, since:

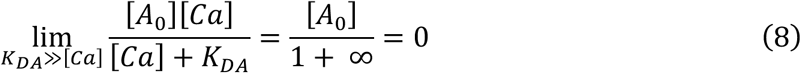

Unless [*A*_0_] is also large, the fraction of calcium that captured by species A will be very low. On the other hand, if A has a very low *K_D_*_A_ ≪ [*Ca*], then

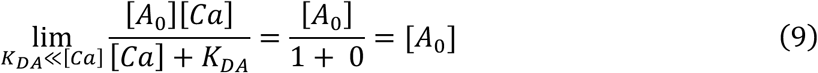

Which indicates almost all of [*A*_0_] will be bound by calcium. The same principle applies to B and C.

This describes the function of each sensor ‘s KD in the partitioning of calcium.

Even though the above analysis is based on the equilibrium state, it illustrates the partitioning calcium based on the dissociation constant and, importantly, it can be applied to a system with coexistence of multiple sensors and buffers.

---

**Figure 2-figure supplement 1.**
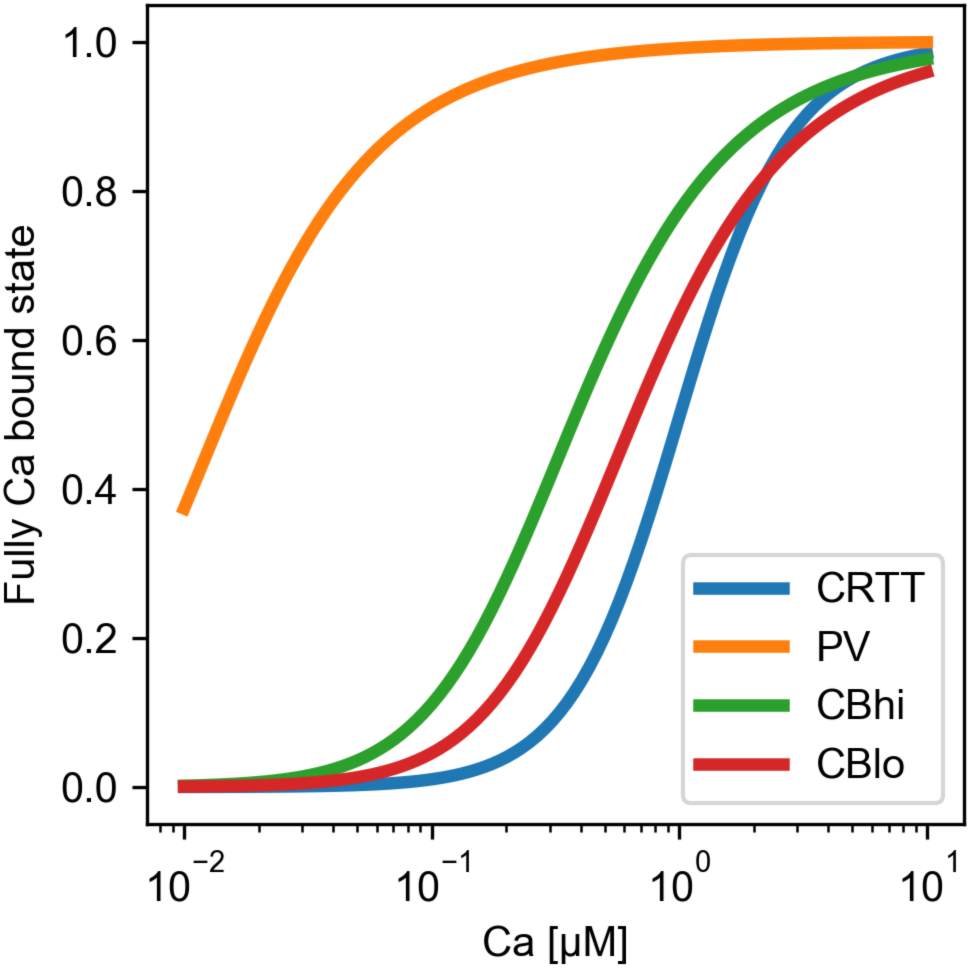
The Calcium binding kinetics for fast buffers of PV, CRTT, and CBhi, CBlo. The dissociation constant KD can be read from the x-axis values at y=0.5. We can see that the KD of all these buffers is lower than that of CaM.

**Figure 4-figure supplement 1.**
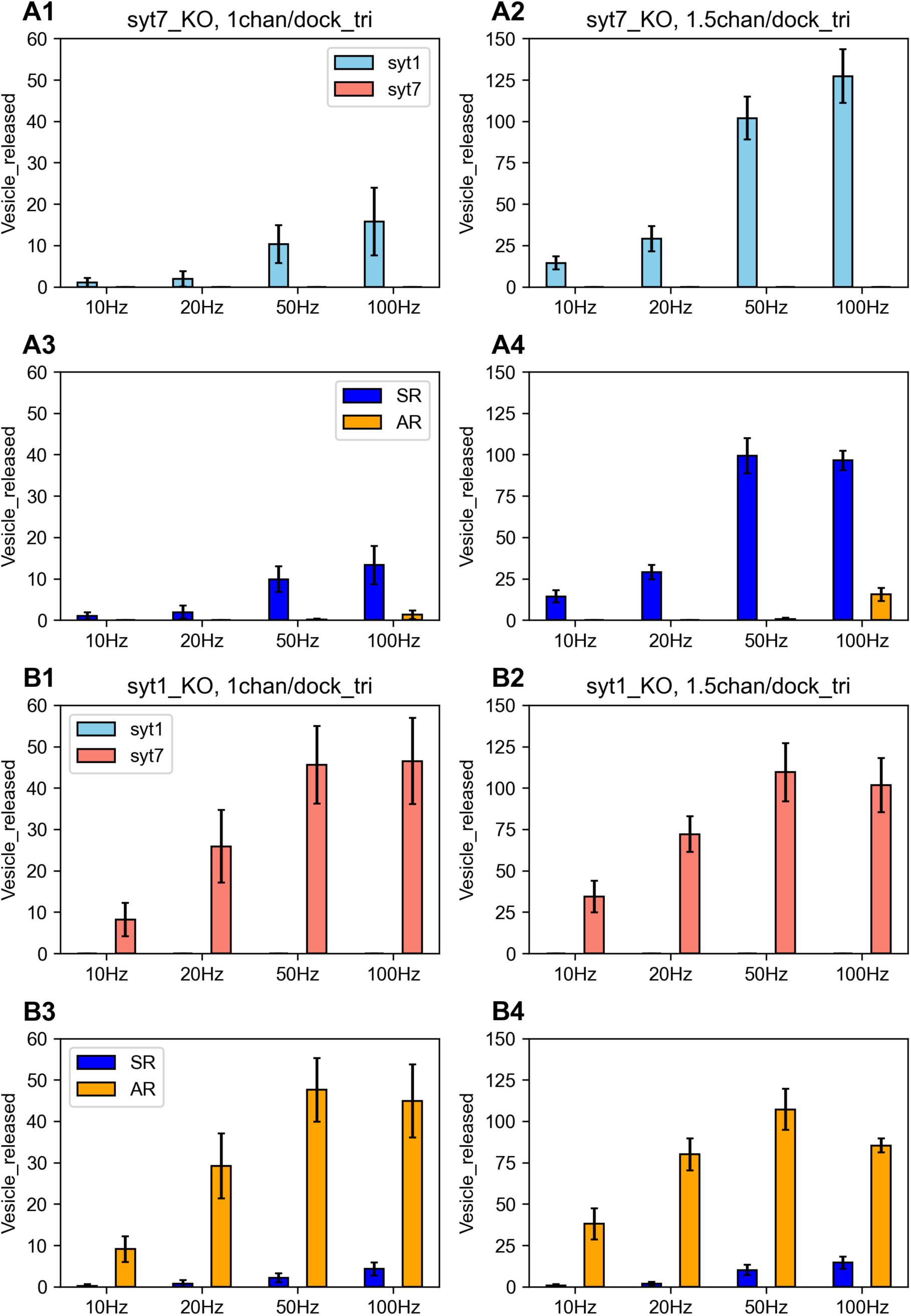
In syt1KO condition syt7 triggered mainly AR; whereas in syt7KO condition syt1 triggered mainly SR. (A1,A2) syt7_KO: vesicle released by syt1; (A3,A4) syt7_KO: vesicle release is charactered by mainly SR. (B1,B2) syt1KO: vesicle released by syt7;(B3,B4): vesicle release is characterized by mainly AR. The error bars are the standard deviations with 10 iterations.

**Figure 6-figure supplement 1.**
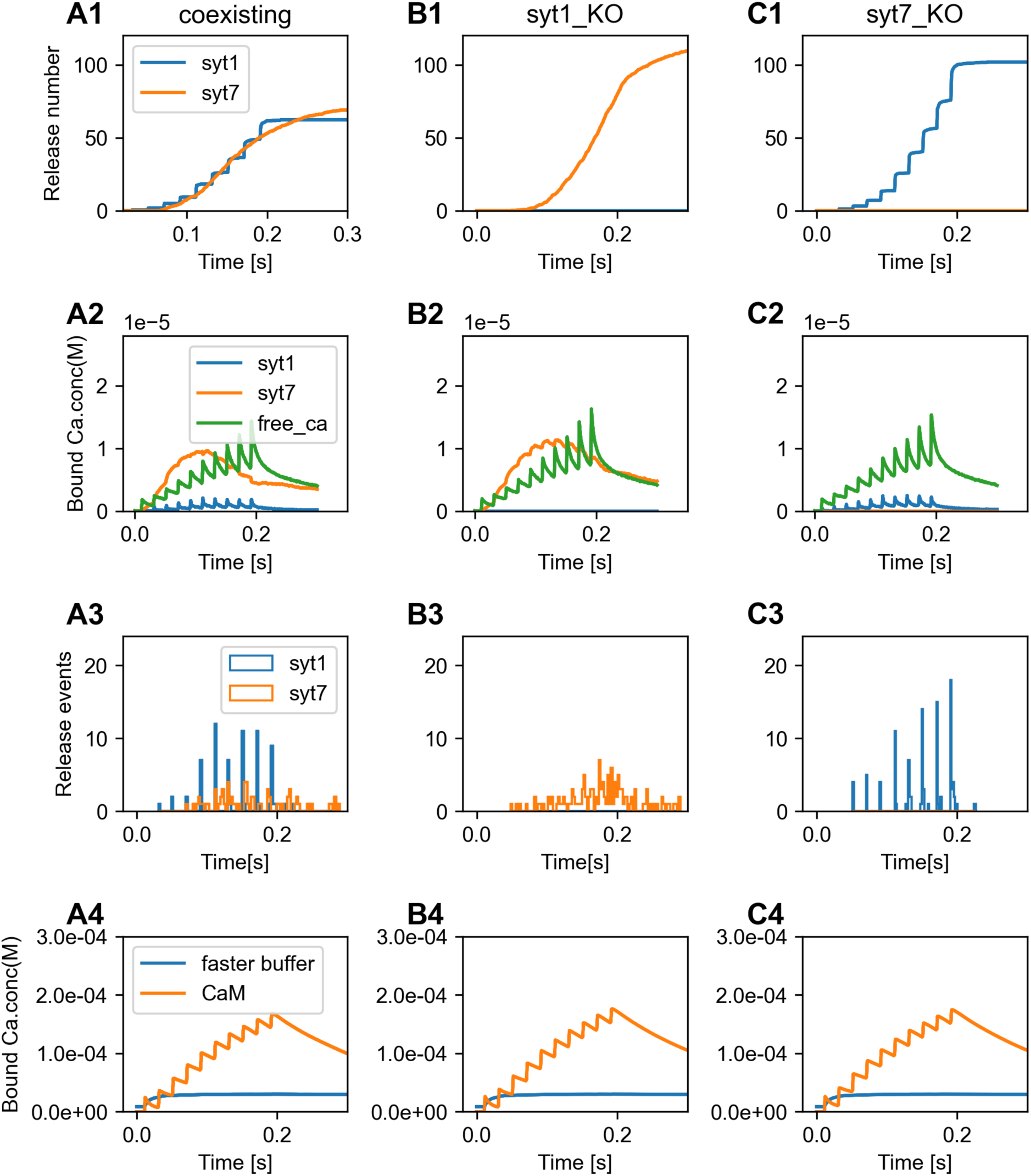
Illustration that free global calcium dynamics cannot predict the release kinetics. Data for 50 Hz stimulation in Fig. 6B1 (1.5 Cachan/dock_tri). (A1-A4): syt1 and syt7 coexist. (B1-B4): syt1_KO. (C1-C4): syt7_KO. In each column with conditions of A,B,C, we show the kinetics of vesicle released by syt1 (blue) and syt7 (orange) in (A1,B1,C1); the concentrations of calcium bound by syt1 (blue) and syt7 (orange) as well as free calcium(green) in (A2,B2,C2); the release events triggered by syt1 (blue) and syt7(orange) in (A3,B3,C3); the concentration of calcium that bound by CaM (orange) and other fast buffers (blue) in (A4,B4,C4). As can be seen in green curve in (A2,B2,C2), the dynamics of free calcium are very similar, but the release pattern are very different. Parameters: syt1 = 4 and syt7 = 4, CaM = 60 µM, Ca_ocon = 2 mM, mean of 10 iterations.

**Figure 7-figure supplement 1.**
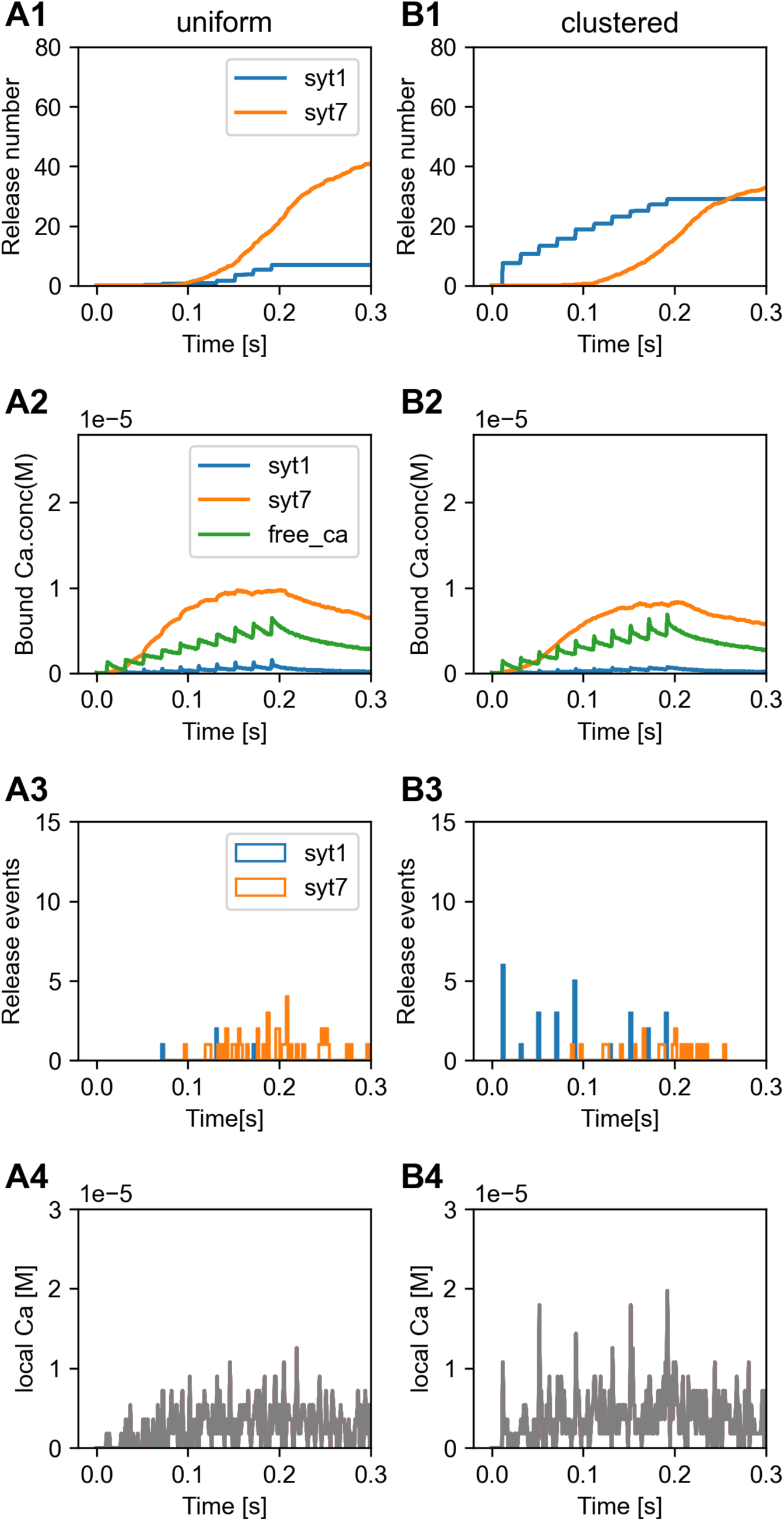
Comparison between uniform distribution and clustered distribution of calcium channels in release kinetics and calcium dynamics shown in Fig. 7 (10 pulses by 50Hz). (A1-A4) uniform distribution: 1 calcium channel in each of 114 docking triangles. (B1-B4): clustered calcium channel distribution, 114 calcium channels in a single docking triangle. (A1,B1) kinetics of vesicle released by syt1 (blue) and syt7 (orange); (A2,B2): the concentrations of calcium bound by syt1 (blue) and syt7 (orange) as well as free calcium(green); (A3,B3) the release events triggered by syt1 (blue) and syt7 (orange). (A4) local concentrations of calcium at tetrahedrons with a calcium channel. (B4) local calcium at the tetrahedron with the clustered channels. Parameters: syt1 = 4 and syt7 = 4, CaM = 60 µM, Ca_ocon = 2 mM, mean of 10 iterations.

**Figure 7-figure supplement 2.**
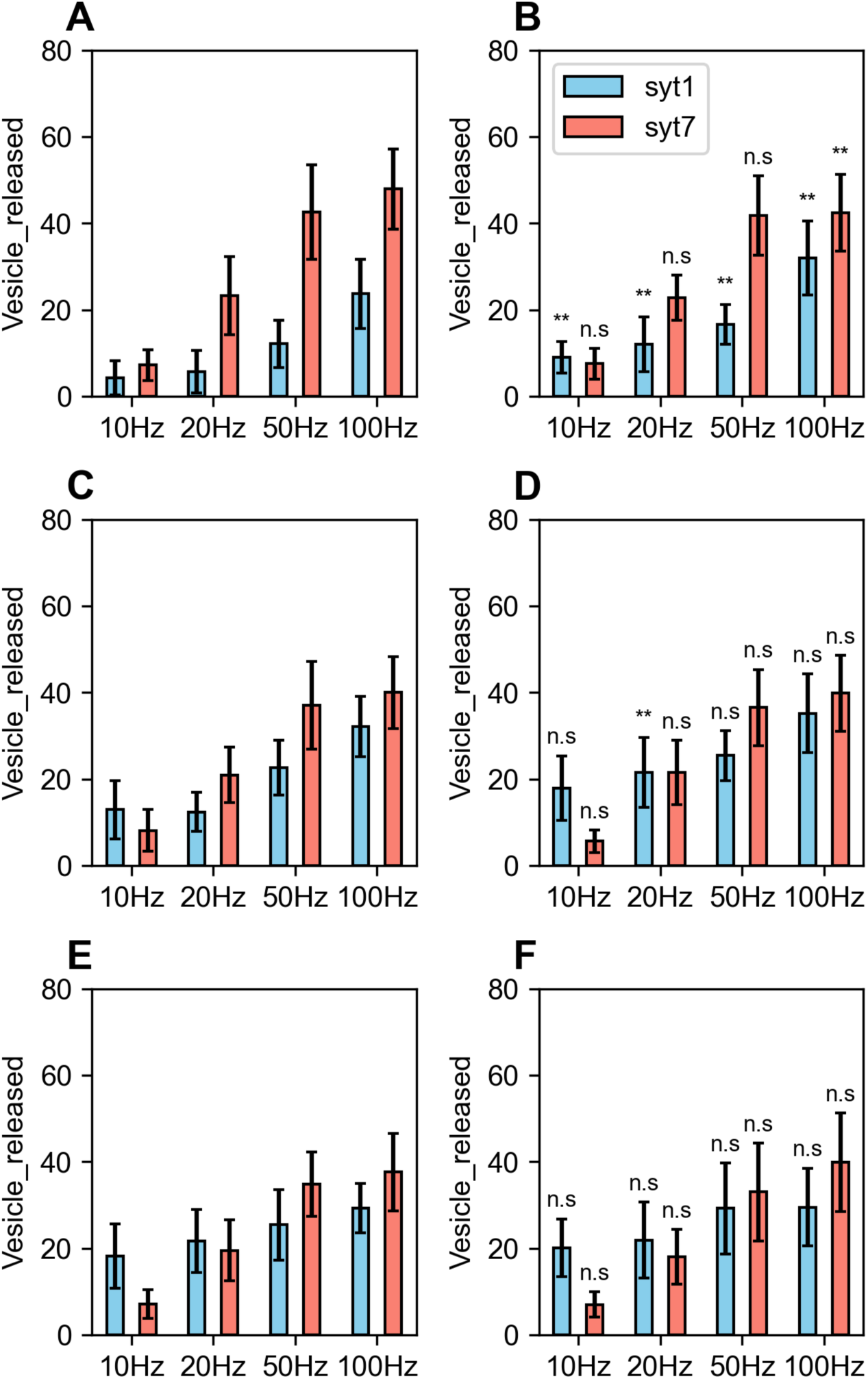
Release patterns with 114 total calcium channel fixed, by varying the clustering pattern by 3, 6, 8, 16, 22, 38 calcium channels in each docking triangle in (A, B, C, D, E, F). For A, B, C: syt1-triggered release is increasing with higher Cachan/dock tri; for D, E, F: both syt1-triggered and syt7 triggered release is saturated, indicating there is a critical value of Cachan/dock_tri above which it will not affect the release patterns. Parameters: syt1 = 4, syt7 = 4, CaM = 60 µM, Ca_ocon = 2 mM, mean of 10 iterations. Statistics: for pair-wise comparison between A and B, C and D, E and F.

**Figure 8-figure supplement 1.**
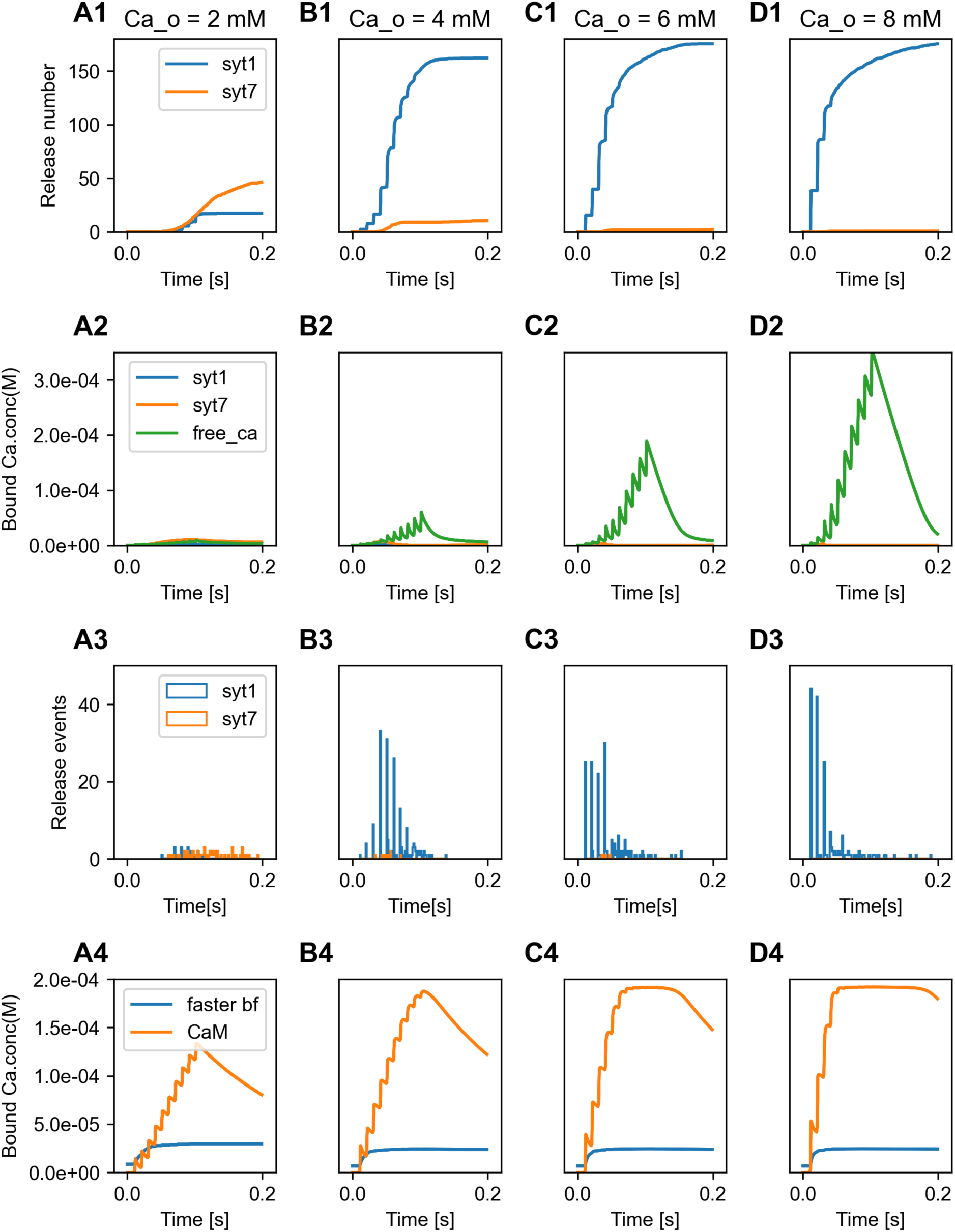
External calcium concentration (Ca_ocon) dependent release by syt1 and syt7. (A) Ca_ocon = 2 mM. (B) Ca_ocon = 4 mM. (C) Ca_ocon = 6 mM. (D) Ca_ocon = 8 mM. For each condition we show on the first row: kinetics of vesicle released by syt1 (blue) and syt7 (orange); on the second row: the concentrations of calcium bound to syt1 (blue) and syt7 (orange) as well as free calcium(green); on the third row: the release events triggered by syt1 (blue) and syt7 (orange); on the last row: the concentration of calcium bound by CaM (orange) and other fast buffers (blue). Parameters: syt1 = 4, syt7 = 4, CaM = 60 µM, mean of 10 iterations.

**Figure 10-figure supplement 1.**
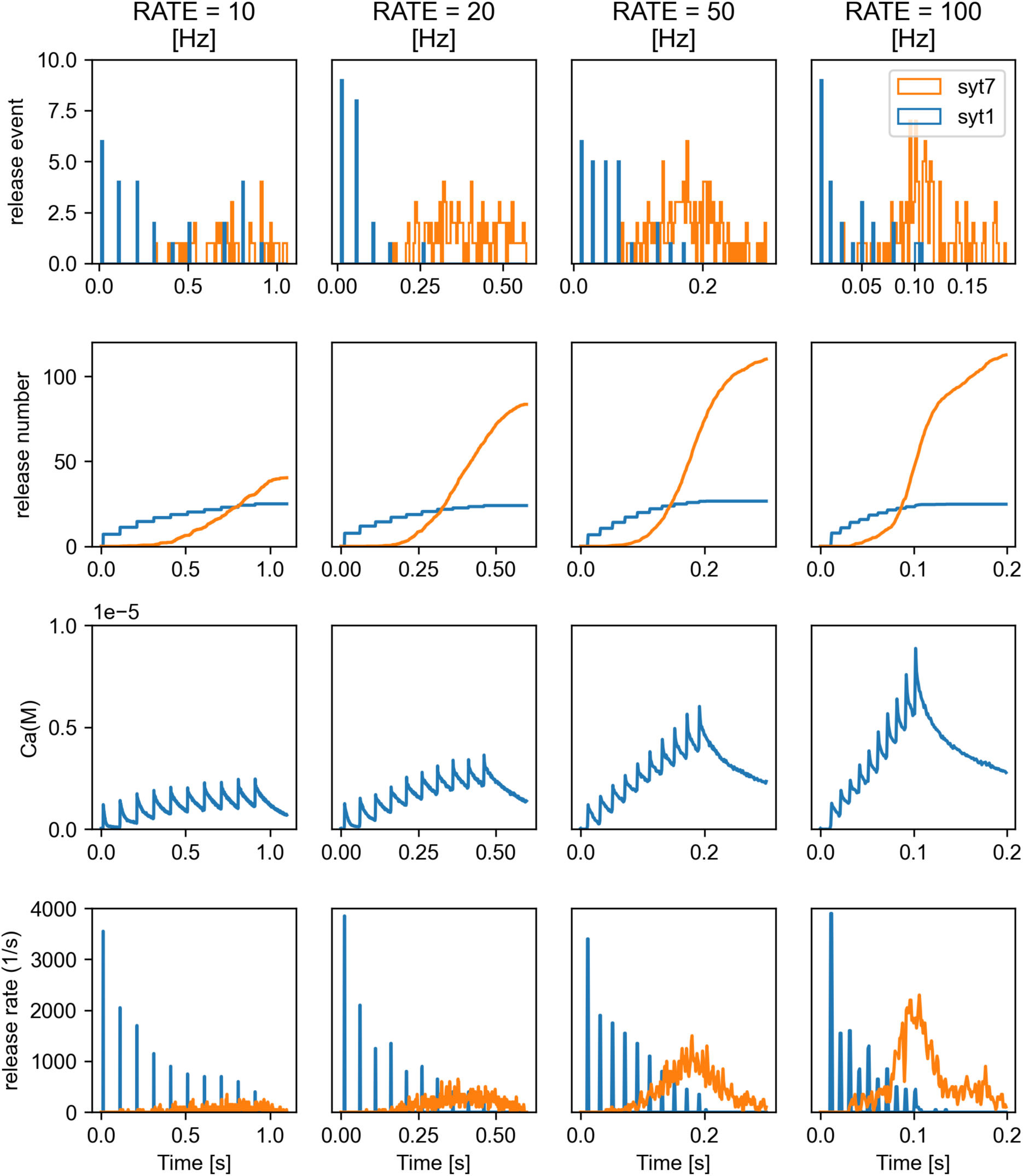
Example illustrating syt7-triggered frequency dependent increase of vesicle release and syt1-triggered frequency independent release. By 10 pulse stimuli at 10,20,50,100Hz, we show first row: vesicle release events triggered by syt1 (blue) and syt7 (orange); second row: vesicle release kinetics triggered by syt1 (blue) and syt7 (orange); third row: free calcium concentration dynamics; last row: releasing rates triggered by syt1(blue) and syt7(orange). Parameters: syt1 = 4 and syt7 = 12, CaM = 60 µM, 114 Ca_chan in 1 dock_tri, Ca_ocon = 2 mM, mean of 10 iterations.

**Figure 10-figure supplement 2.**
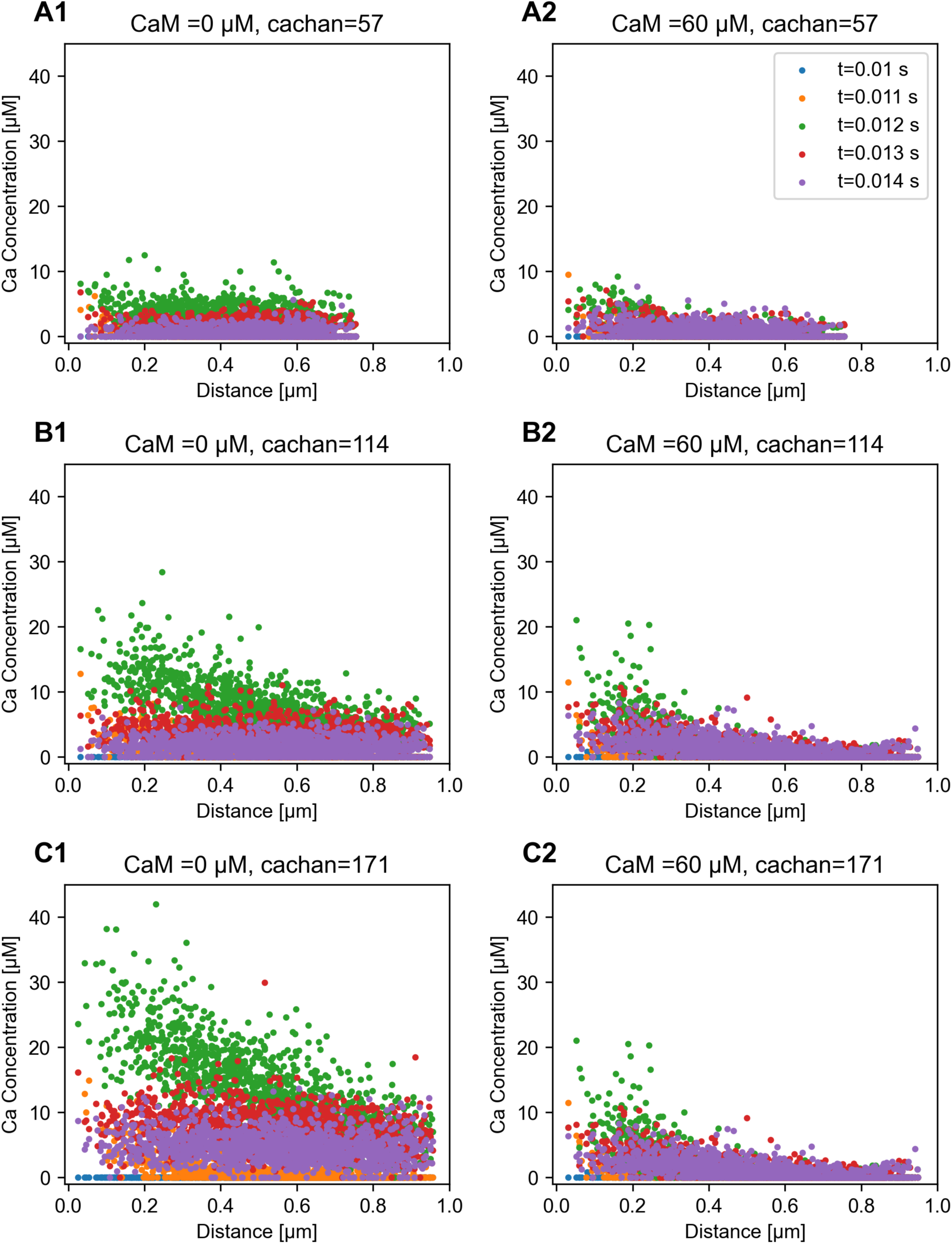
Increase of the free calcium concentration with increasing calcium channel numbers in a single cluster. (A) 57. (B) 114. (C) 171 channels in the cluster. CaM = 0 µM (left) and CaM = 60 µM (right). This figure shows the effect of CaM on calcium buffering which is at a time scale of sub-millisecons and spatial scale of around 0.3 µm. Other parameters: ca_ocon = 2 mM, 10 pulses stimuli with100 Hz: at t = 0.011 s the voltage is clamped to 40 mV, at t = 0.012 s, the voltage switches back to -61 mV. The location of the docking triangles for placing the calcium channel clusters are randeomly selected.

**Figure 10-figure supplement 3.**
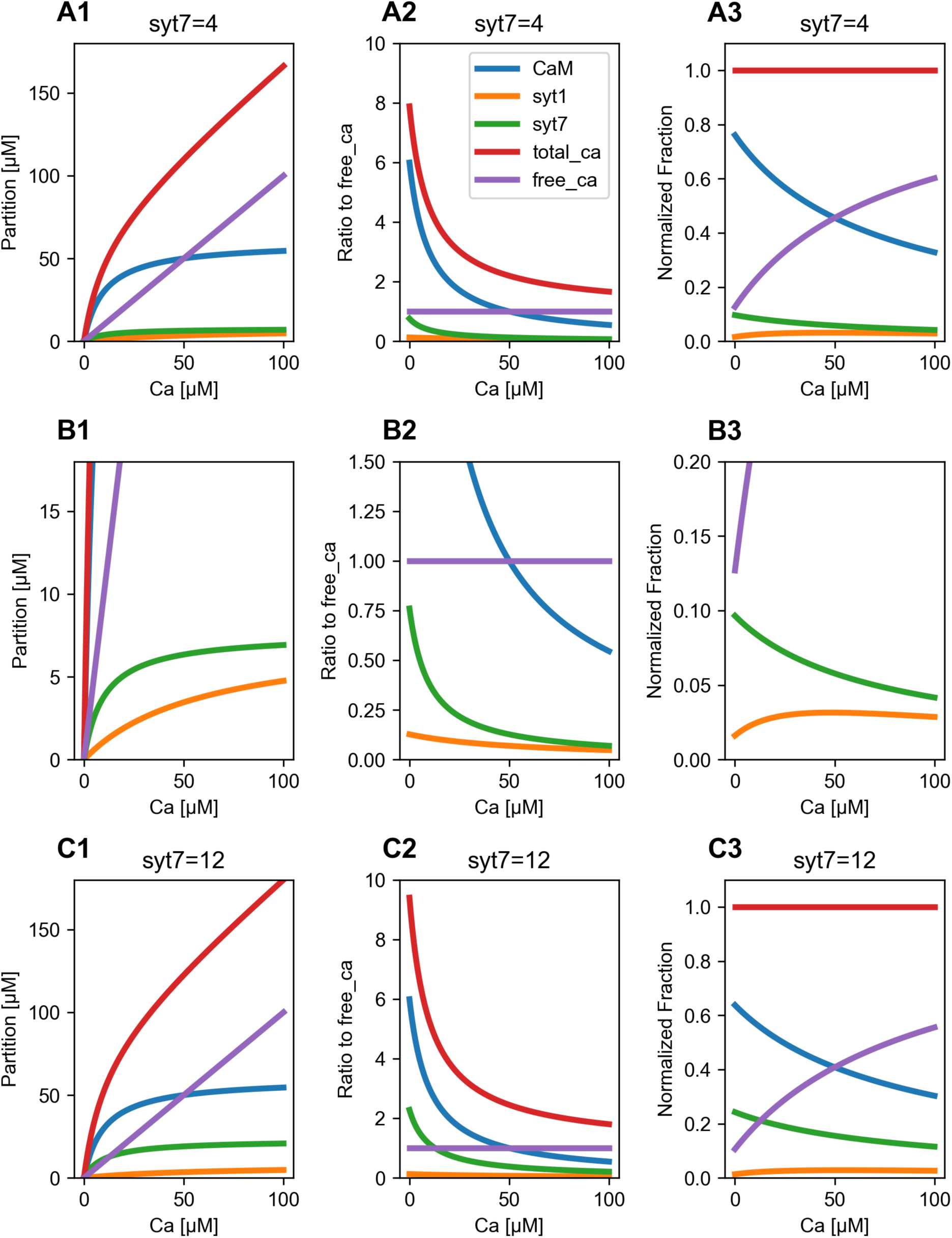
Calcium partitioning for different free calcium levels with (A1-A3, B1-B3) syt1= syt7 = 4 and (C1-C3) syt1 = 4 and syt7 = 12 . (A1) free calcium concentration (purple), partition of CaM_bound (blue), partition of syt1_bound (orange), partition of syt7_bound (green) and total calcium (red); (A2) the ratios of these quantities relative to free calcium. (A3) the fraction of these quantities relative to total calicum. (B1-B3) zoomed in for (A1-A3). Parameters: syt1 = 300 vesicles * 4/vesicle ∼ 7.6 µM, CaM KD and syt7 KD are both 10 µM, and syt1 KD is 60 µM.

**Table S1.** Kinetic rates for the other buffers used in the model.

| Buffer | On rate[$M^{-1}s^{-1}$)] | Off rate [$s^{-1}$] |
| --- | --- | --- |
| PV | 1.07e8 | 0.95 |
| CBhi | 1.1e7 | 2.607 |
| CBlo | 8.7e7 | 35.76 |
| CRTT | 3.6e6 | 53 |
|  | 3.1e8 | 20 |

## Notes

### Competing Interest Statement

The authors have declared no competing interest.

